# Whole-Brain, Region-Specific Astrocyte Reactivity and Morphological Remodeling After Diffuse Traumatic Brain Injury in A Gyrencephalic Ferret Model

**DOI:** 10.64898/2026.07.16.739056

**Authors:** Amirhossein Bagherian, Camila Perez, Allison Kosub, Rianna Chalijah Ysabelle Gonzales, Alexander Patterson, Kevin F. Bieniek, Morteza Seidi, Marzieh Memar

**Affiliations:** Department of Biomedical Engineering and Chemical Engineering, The University of Texas at San Antonio, TX, USA; Department of Pathology and Laboratory Medicine, The University of Texas at San Antonio, TX, USA; Glenn Biggs Institute for Alzheimer’s & Neurodegenerative Diseases, The University of Texas at San Antonio, TX, USA; Department of Mechanical Engineering, The University of Texas at San Antonio, TX, USA

**Keywords:** Traumatic brain injury, GFAP, Diffuse axonal injury, CHIMERA, Deep learning segmentation, Ferret model, Brain atlas mapping, Astrocyte morphology

## Abstract

Traumatic brain injury (TBI) triggers pathological cascades that evolve across acute, subacute, and chronic phases. Astrocytes play a central role across these phases, and astrocyte reactivity is commonly evaluated using glial fibrillary acidic protein (GFAP) immunolabeling. However, in many TBI studies GFAP changes are characterized qualitatively or with manual or simple threshold-based measures on a small set of sections, limiting throughput and constraining analysis of region-specific heterogeneity in astrocyte responses. To overcome these limitations, we employed a ferret model of diffuse TBI (5 TBI, 5 sham), leveraging the ferret’s gyrencephalic cortex, human-like regional fractional brain volumes, and astrocyte features that more closely resemble the human brain than rodent models. An AI-driven segmentation model validated for GFAP-stained ferret histology was integrated with atlas-based mapping to achieve whole-brain, region-resolved quantification of astrocyte reactivity over an average of 10 coronal slices per animal. Morphometric analysis using a custom SMorph-based pipeline characterized branching complexity and spatial domain features across defined regions. At seven days post-injury, TBI animals showed elevated astrocyte reactivity and hypertrophic remodeling, with significant expansion of convex hull area and elongation of secondary branches at the whole-brain level, most pronounced in the atlas-defined gray-matter region and cerebellum and brain-stem subregions, whereas white-matter showed a similar but less marked trend. Morphological changes were also detected in the hippocampus that did not show significant increases in astrocyte reactivity, indicating that structural remodeling represents a partially independent dimension of the astroglial response. These regional patterns are consistent with expected large tissue deformation and axonal strain in brainstem-cerebellar pathways and gray-matter at gray-white junctions in sagittal rotation, motivating future computational studies to quantify these links more directly. By combining region-resolved GFAP mapping with large-scale morphometry, this work provides a scalable framework for region-specific astrocyte mapping to support future multimodal, computational, and targeted neuroprotective studies.

## Introduction

Traumatic brain injury (TBI) represents a significant public health concern, affecting millions of individuals each year and ranging from mild concussions to severe brain damage. According to the Centers for Disease Control and Prevention (CDC), approximately 586 TBI-related hospitalizations and 190 TBI-related deaths occur daily across the United States. These injuries arise from falls, vehicle accidents, and sports-related activities, often involving external forces or rapid head rotation, with contact-sports alone accounting for about 10% of all reported TBIs [6, 17, 38, 86]. Depending on mechanism, severity, and frequency of exposure, TBIs can lead to short-term effects, including altered mental state, dizziness, and speech or sensory difficulties, as well as long-term outcomes such as behavioral and emotional changes and elevated risks of neurodegenerative disorders, anxiety, and depression [41, 91].

Despite this burden, the pathophysiology of the brain’s acute and chronic responses to TBI remain incompletely understood. TBI initiates cascades that disrupt neurons, glia, vasculature, and the blood–brain barrier, driving acute damage and long-term neuroinflammatory responses [75]. These neuroinflammatory cascades involve coordinated responses across multiple cellular populations, including microglia, astrocytes, neurons, and endothelial cells. Quantitative analysis of neuropathology can therefore provide insight into cell-type- and region-specific responses as well as help inform therapeutic strategies [26]. Within this neuroinflammatory response, one key hallmark is astrocyte reactivity, commonly assessed by upregulation of glial fibrillary acidic protein (GFAP), which serves as a marker of astrocyte injury and reactive cytoskeletal remodeling in response to trauma and other central nervous system insults [80]. Although GFAP captures only a subset of reactive astrocyte changes, it is one of the most extensively studied astrocyte markers and is widely used in both preclinical and clinical TBI research: in animal models as a histopathological readout and as a biofluid biomarker, and in patients primarily as a blood- and CSF-based biomarker [30, 44, 47, 71, 84, 93], enabling direct comparison between tissue-level pathology and circulating injury signals. Detailed quantification of astrocyte reactivity throughout the brain following TBI can provide insight into how injury mechanism and severity relate to the whole-brain, regional, and tissue-level distribution of damage and can establish a foundation for targeted and region-specific assessment of therapeutic interventions [68].

Astrocytes play a pivotal role in the brain’s response to neurotrauma due to their involvement in blood–brain barrier maintenance, homeostasis regulation, metabolic support, and neurotransmission [67, 109]. Astrocytes guide axons, form synapses, refine neuronal networks, and stabilize the central nervous system by buffering ions, regulating pH, and detoxifying ammonia, while also influencing circuit function by shaping synapse dynamics and neurotransmitter availability [12, 46, 58]. Following injury, astrocytes respond rapidly and remain reactive for extended periods, undergoing hypertrophy, proliferation, morphological changes, and glial scar formation that limits tissue damage and promotes repair [14, 15, 66, 118]. Their abundance, functional diversity, and involvement in both short- and long-term outcomes make them valuable markers for studying TBI pathology, severity, and progression [113].

In the broader context of TBI, neuroinflammatory cascades persist well beyond the acute phase, driven by sustained glial activation and ongoing tissue remodeling [65]. In contrast, acute axonal injury markers fluctuate and decline over time, limiting their utility for monitoring subacute and chronic pathology [65], whereas astrocyte reactivity persists for extended periods and provides an opportunity to track ongoing injury-related inflammatory and remodeling processes [15]. Prolonged astrogliosis contributes to chronic neurodegeneration and long-term cognitive and behavioral impairments, making GFAP-based astrocyte reactivity a particularly valuable histopathological marker for linking diffuse brain injury to its functional consequences [31, 50, 85].

Moreover, astrocyte abundance, morphology, and function are not uniform across the brain and vary by region, with protoplasmic astrocytes predominating in gray matter and fibrous astrocytes in white matter [113]. These subtypes differ in their morphological structures including territorial organization, cytoskeletal architecture, and interactions with vasculature, neurons, and synapses, which lead to region-specific patterns of GFAP expression and process morphology at baseline. Consequently, astrocyte reactivity after injury is likely region-dependent, such that diffuse TBI may result in region-specific structural responses, for example distinct astrocyte responses in cortical gray matter compared with long-range white-matter tracts. Capturing this heterogeneity requires region-resolved analysis of astrocyte reactivity and morphology rather than bulk measures of GFAP signal intensity from a limited number of sections. However, truly systematic, spatial analysis of astrocyte reactivity and morphology remains limited in the current literature and demands controlled injury severity and timing, standardized tissue processing, and broad anatomical sampling, conditions that are rarely achievable in human clinical TBI studies. These limitations necessitate animal models that recapitulate key features of human TBI pathology, including mixed gray–white matter involvement and gyrencephalic cortical architecture, and provide a controlled platform for systematically evaluating regional astrocyte responses.

To meaningfully study astrocyte responses to TBI, animal models must capture critical aspects of human neuroanatomy and biomechanics. Rodent models (mice, rats) are widely used to uncover TBI mechanisms and test treatments [21, 56, 74, 79, 100]; however, their translational relevance is limited by the lissencephalic brain structure and relatively low white-matter content of rodent brains. Ferrets, the smallest gyrencephalic species, offer a more human-like neuroanatomical framework, with greater similarity in hippocampal positioning, cortical folding, and white-to-gray matter ratios [95, 117], while maintaining lower costs than larger gyrencephalic models such as pigs. Importantly, ferret astrocytes also exhibit several morphological, transcriptional, and functional characteristics that are homologous to human astrocytes than rodent astrocytes, namely longer and more highly branched processes, greater proliferative capacity, distinct calcium signaling responses, and expression of several astrocyte-specific genes shared with humans, supporting their utility for studying astrocyte responses to TBI in higher-order brains [89]. In conjunction with the most appropriate species selection, experimental injury paradigms range from focal cortical impact to diffuse axonal injury approaches, such as rotational acceleration, which more closely replicate human TBI biomechanics [62, 81].

While animal models enable comprehensive evaluation of astrocyte responses, most pathological analysis in animal studies of TBI still rely on manual or threshold-based cell classification and quantification, which is limited in scope, prone to variability, and often restricted to a small number of slices or regions, potentially missing substantial astrocyte responses elsewhere [90, 94]. This limitation is particularly relevant in diffuse TBI, where injury may span multiple, spatially distinct regions [25, 28, 94, 113]. Recent advances in artificial intelligence (AI) now enable automated, high-throughput object classification and segmentation of astrocyte reactivity, expanding the capacity to map cells and pathology on a system-wide scale, offering region-resolved quantitative insight not achievable with conventional manual approaches. However, astrocyte reactivity is not only reflected by changes in GFAP expression, but also by structural remodeling of astrocyte processes, which is often described qualitatively rather than quantified systematically throughout the brain. While regional GFAP-immunoreactive area coverage provides an important region-resolved proxy of adaptive astroglial responses to TBI, segmentation alone cannot fully capture the structural complexity of astrocyte remodeling. Systematic morphological analysis enables quantification of astrocyte process architecture from segmented masks, including branching complexity and ramification metrics, across anatomically mapped brain regions. Although computational tools for quantitative cell-morphology analysis are emerging, comprehensive and user-friendly software pipelines remain limited, particularly for glial cells. Existing approaches include open-source workflows (e.g., FIJI/ImageJ with Simple Neurite Tracer) [107], glia-focused toolkits such as GliaMorph [64], and commercial 3D reconstruction platforms such as Neurolucida [98], all of which have been adapted for astrocyte analysis in specific contexts. Among these, SMorph is one of the few pipelines developed explicitly for automated astrocyte morphological analysis [96].

Building on considerations of species selection, injury paradigm, and the need for comprehensive region-specific analysis, we employed a ferret model of TBI using Closed-Head Impact Model of Engineered Rotational Acceleration (CHIMERA) injury exposure, which applies rapid rotational forces to produce diffuse TBI. To quantify TBI-induced astrocyte reactivity, we used a deep-learning model to segment reactive astrocytes and an automated mapping algorithm to assign segmented cells to regions in a ferret brain atlas, achieving high-resolution, region-specific, whole-brain mapping that overcomes the common limitation of targeted cell profiling. In addition, we incorporated large-scale morphometric analysis of segmented astrocytes using a custom SMorph-based pipeline to quantify astrocyte process architecture across anatomically mapped brain regions. This integrative framework minimizes observer bias, expands the spatial scope of analysis, and enables detection of subtle regional differences in astrocyte reactivity, offering novel region-specific insights into astrocyte responses after diffuse TBI that may inform future prognostic and targeted therapeutic intervention. This atlas-registered, quantitative framework provides a foundation for identifying brain regions that may exhibit disproportionate astroglial remodeling after diffuse injury and for guiding future region-specific therapeutic studies using the ferret TBI model.

## Materials and Methods

### Human Relevance of The Ferret TBI Model for Regional Astrocyte Analysis

As noted above, prior work has shown that ferret astrocytes share key morphological features with human astrocytes [89], supporting their use as a translational model for astrocyte-focused TBI studies. However, to further evaluate the human relevance of the ferret TBI model for region-specific pathological quantification, we calculated regional fractional volumes across commonly used preclinical TBI species and compared them with regional fractional volume in humans. For each species, regional fractional volume was computed by dividing the volume of each brain region by the total brain volume of that species. Using established MRI atlases from humans (mixed-sex population template; 609 males, 721 females) [51, 88] rhesus macaques (10 adults, 2–11 years; 7 males, 3 females)[16], pigs (20 males, 4 weeks) [35], ferrets (26 *in vivo* and 8 *ex vivo* scans, 5–10 months)[49], rats (6 adult male Wistar) [53], and mice (14 males, 8 weeks old) [52, 110, 111], we derived regional fractional volumes across these species (Figure 1 and Supplementary Table 1).

**Figure 1.**
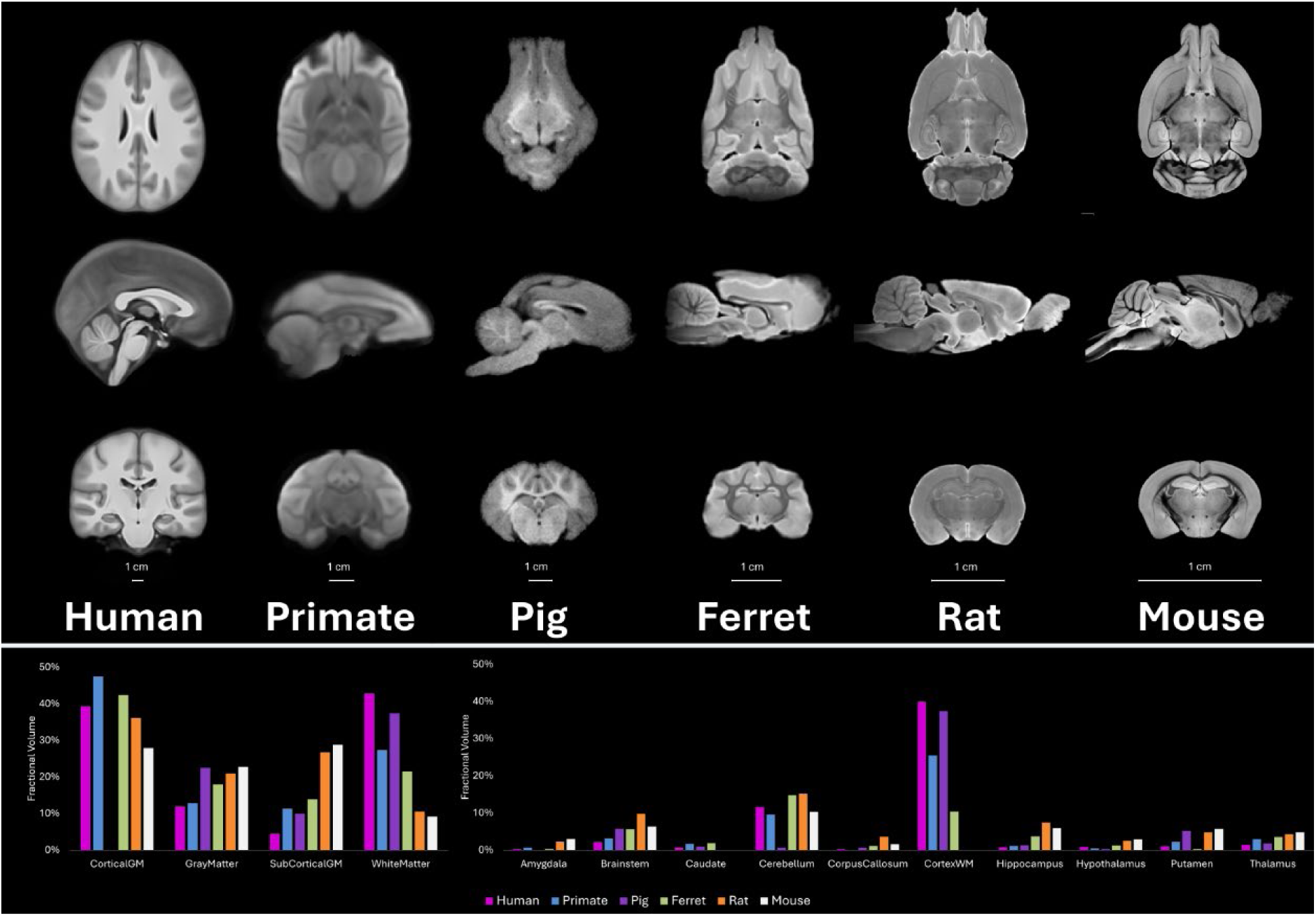
Top: Cross-species MRI templates illustrating macroscopic neuroanatomical features of the human, non-human primate, pig, ferret, rat, and mouse brain. Bottom: Regional fractional volume analysis of key neuroanatomical regions across species.

### Animal Experiments

All animal procedures were approved by the Institute of Animal Care and Use Committee of the University of Texas at San Antonio (IACUC #MF001) and conducted in accordance with National Institutes of Health guidelines. A total of 10 male ferrets (*Mustela putorius furo*), approximately 6 months of age (equivalent to ∼18 human years), were randomly assigned to TBI and sham groups to investigate the astroglial response of diffuse TBI. Ferrets were purchased from Marshall Bioresources (North Rose, NY, USA) and housed 3–4 per cage under a 12-h light/12-h dark cycle, with *ad libitum* access to food and water. Animals were acclimated for approximately 10 days before experimentation and monitored daily throughout the study.

On the day of experiment, prior to anesthesia induction, ferrets were fasted for 1–2 hours to reduce anesthesia-related side effects. Animals were then injected intramuscularly with Acepromazine (SKU 003845, Covetrus, Portland, ME, USA) at 0.1 mL/kg body weight. The stock Acepromazine solution (10 mg/mL) was diluted 1:10 in sterile saline and administered 15 minutes prior to anesthesia to facilitate a smooth recovery. Anesthesia was induced in a chamber with 5% isoflurane in oxygen at 4 liters per minute (LPM). Depth of anesthesia was verified by the absence of a rear-toe-pinch reflex, after which ferrets were transferred to a surgical table equipped with a water-circulating heating pad set at 100 °F (38 °C) for thermal support. Anesthesia was maintained via a nose cone at 1.5–2.5% isoflurane in 2 LPM oxygen, and the rear-toe-pinch reflex was checked periodically. After induction, ferrets received 10 mL of saline subcutaneously to counteract potential effects of prolonged anesthesia and blood collection. Buprenorphine SR (BUPREN-INJ008VC, Wedgewood Pharmacy, Swedesboro, NJ, USA) was administered at 0.012 mL/kg body weight for analgesia. Throughout anesthesia, vital signs, including heart rate (180–250 beats/min), oxygen saturation (95–100%), and rectal temperature (100–104 °F) [87], were monitored every 15 minutes until full recovery.

For the TBI group (n = 5), animals were briefly disconnected from anesthesia, placed in a supine position on the padded interface of the CHIMERA system (Figure 2, Vancouver, Canada), and secured using two Velcro straps at the shoulders and abdomen. Animals then received a single head impact at ∼16 m/s (25.6 J), delivered by an air-driven actuator set at 60 psi. A helmet-like interface prevented direct rod contact with the skull, reducing the risk of fractures. The impact caused rapid head rotation around the fixed shoulder restraint, creating inertial forces known to induce diffuse TBI. Immediately after impact, animals were returned to the surgical table and reconnected to anesthesia. The ferret CHIMERA model has been shown to effectively reproduce rapid rotational head injury with outcomes resembling human TBI [48, 62, 63], and similar impact parameters (∼26–27 J, 60–65 psi) have been reported to induce diffuse axonal injury [62]. The sham group (n = 5) underwent identical anesthesia and handling without head impact. After completion of procedures, isoflurane was discontinued while oxygen at 2 LPM was maintained until the animals regained consciousness; they were observed for 15–30 minutes post-recovery, or longer if needed until they returned to normal activity before being returned to their home cages.

**Figure 2.**
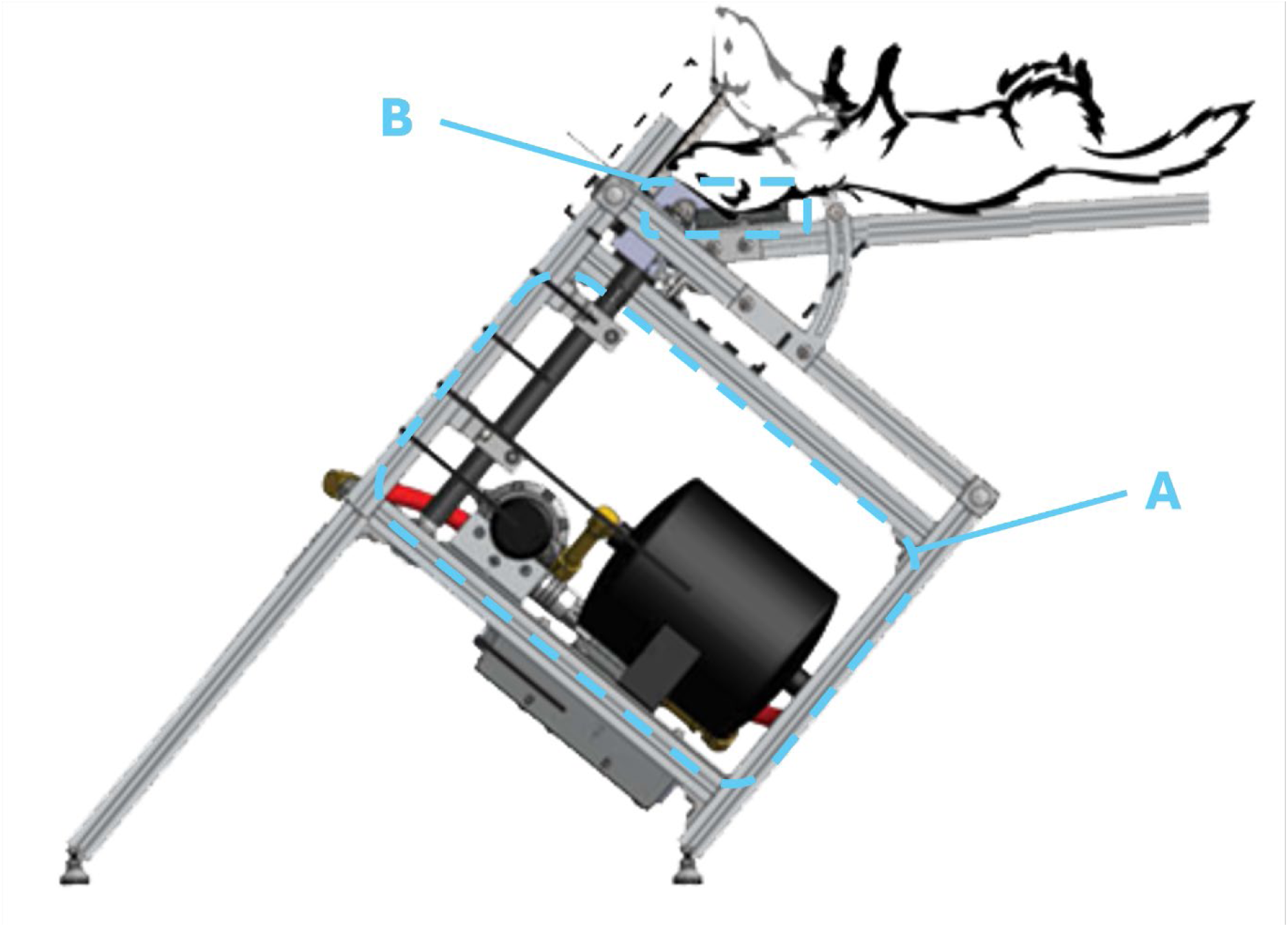
Schematic of the CHIMERA device (Vancouver, Canada) used to deliver rapid rotational head impacts in ferrets. The system is equipped with an air-driven actuator (A) and a padded head interface (B), producing controlled inertial rotation suitable for modeling diffuse traumatic brain injury.

Animals were euthanized 7 days post-injury using a pentobarbital-based euthanasia solution (Virbac, Westlake, TX), administered via intracardiac injection at 1 mL. Prior to euthanasia, 1 mL of heparin (10,000 units/mL; Covetrus, Portland, ME, USA) was administered via intracardiac injection to prevent clot formation. Perfusion fixation was performed immediately post-euthanasia using an automated pressure perfusion system (Leica Biosystems, Nussloch, Germany). First, 1 L of saline was perfused to clear the vasculature, followed by 1 L of 4% paraformaldehyde (PFA) in phosphate-buffered saline (PBS) for tissue fixation.

### Brain Tissue Processing, Slicing, and Paraffin Embedding

Following perfusion, brains were carefully extracted and post-fixed in 4% PFA in PBS for 24– 48 hours at 4 °C (Figure 3A). Brains were then sectioned in the coronal plane at a uniform thickness of 2 mm (Figure 3B) using microtome blades and a custom-made adjustable ferret brain slicer. Each 2 mm slice was placed in an individual cassette, incubated in 4% PFA in PBS for 4 hours at 4 °C, rinsed twice with PBS, and stored in 70% ethanol in PBS at 4 °C until processing by the UT San Antonio Pathology Core. Brain slices were processed using a standard automated tissue processor, which sequentially dehydrated the tissue with graded ethanol, cleared it with xylene, and infiltrated it with molten paraffin. Following processing, slices were embedded in paraffin using the Tissue-Tek system (Sakura Finetek, USA) to ensure long-term preservation, structural stability, and compatibility with precise microtome sectioning and histological staining.

**Figure 3.**
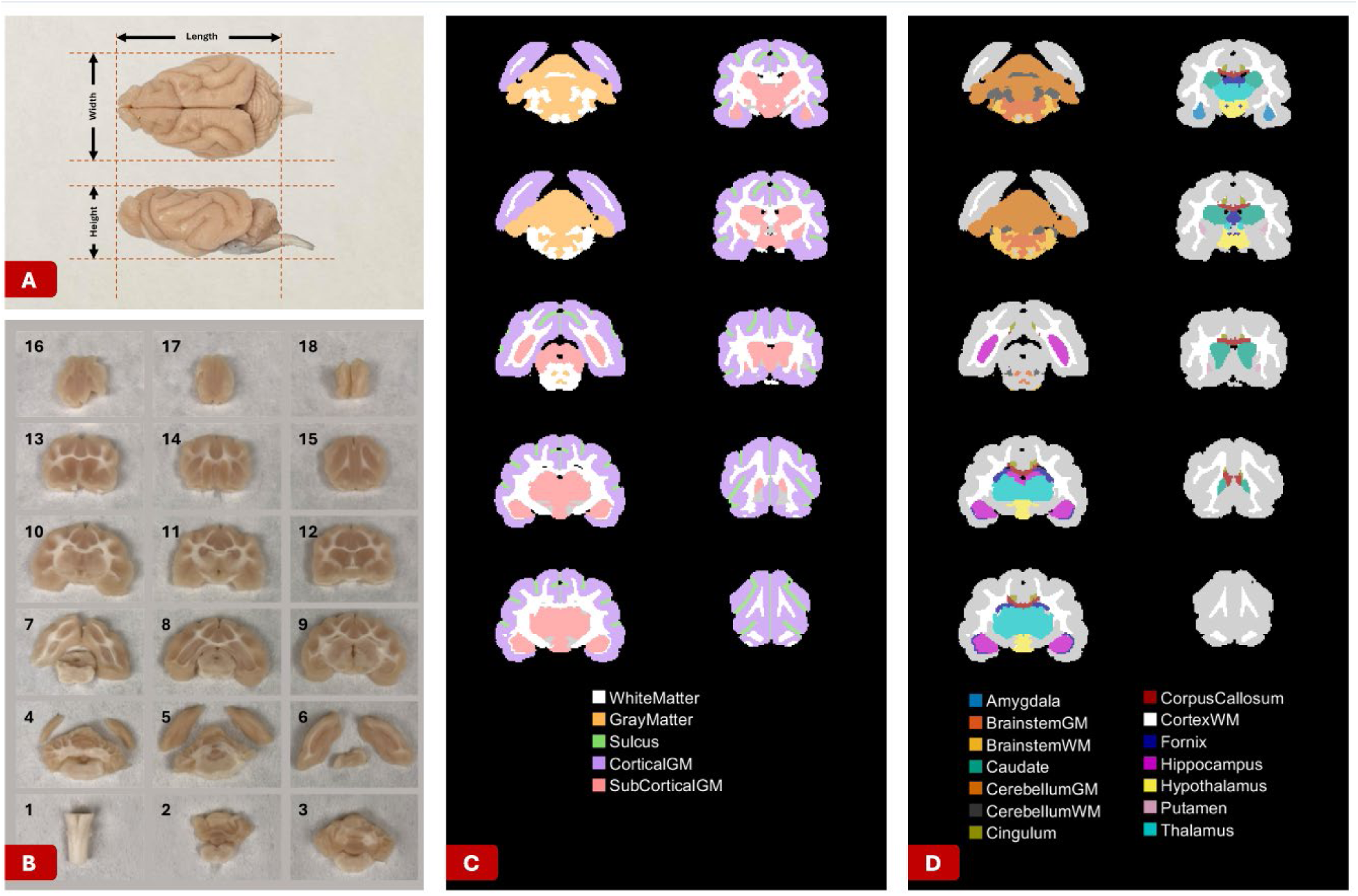
A) Dimensions of the perfused ferret brain measured prior to sectioning. B) Coronal sections obtained at 2-mm intervals using a custom ferret brain slicer. C) Ferret brain atlas illustrating main regions across representative anterior–posterior slices. D) Corresponding atlas subregions detailing specific neuroanatomical regions used for regional quantification of astrocyte reactivity.

### Immunofluorescent Staining and Imaging

Following paraffin embedding, one 6–10 μm tissue section was cut from each 2 mm paraffin block using a microtome, transferred to a 38 °C water bath, mounted on microscope slides (22037246, Fisher Scientific), and allowed to adhere overnight at room temperature (22 °C). Rehydration was performed with two xylene washes, three graded ethanol washes (100%, 95%, 70%; Flex-100 diluted with DI water), and a final 1X Tris-buffered saline (TBS, pH 7.6) wash. Antigen retrieval was carried out in citrate buffer (pH 6.0) using a pressure cooker for 15 minutes at high pressure, followed by three 10-minute washes in TBST (1X TBS with 0.1% Tween-20, pH 7.4). For immunostaining, slides were blocked with 5% bovine serum albumin (BSA) in 1X TBS for 1 hour at room temperature, then incubated overnight at 4 °C with primary antibody anti-GFAP (ab4674, chicken polyclonal, Abcam) diluted 1:500 in blocking buffer. After three 10-minute TBST washes, slides were incubated for 1 hour at room temperature with FITC-conjugated donkey anti-chicken IgY (Invitrogen, ThermoFisher) diluted 1:200 in blocking buffer, washed three additional times in TBST, and mounted with VectaShield Antifade (Vector Laboratories) under coverslips.

Slides were imaged within 1 week using a Keyence BZ-X800 fluorescence microscope (Keyence Corporation of America, Elmwood Park, NJ) at 20× magnification. Exposure times ranged from 1/15 s to 1/5 s with excitation intensity at 40%. An autofocus capture with an 11-slice z-stack was used to generate a single focused composite image for each field of view, maintaining consistent resolution across the tissue depth, and images were saved in 48-bit format for subsequent analysis.

### Pathology Quantification, Mapping, and Regional Analysis

To comprehensively quantify astrocyte reactivity across the ferret brain, an in-house deep learning model purpose-built for automated segmentation of astrocytes in ferret pathology images was employed [1, 7]. We implemented a semi-automatic analysis pipeline in which pathology images were overlaid onto a 3D MRI-based ferret brain atlas [49], constructed from five- to ten-month-old animals, similar in age to the ferrets used in this study, to determine both the extent and anatomical location of astrocyte reactivity and remodeling in distinct regions. The dataset comprised 97 pathology slides, averaging 10 slices per animal, all processed with consistent staining, imaging, and analysis protocols to enable rigorous whole-brain quantification.

Pre-processing combined manual and semi-automatic steps. Despite standardized acquisition and preparation, all images underwent intensity normalization to correct inter-slide variability: for each slide, mean pixel intensity was adjusted to the dataset-wide average. Alignment was verified visually; non-horizontal slides were rotated in 1° increments from −25° to +25°, with optimal orientation selected manually, and tissue boundaries were delineated by hand to exclude background.

Microscopy images were then subdivided into 631×486 pixel patches. The previously developed astrocyte segmentation model for ferret brain images, based on a U-Net++ architecture with a VGG19 backbone and trained on expert-annotated images spanning diverse astrocyte reactivity states [7], was applied to the entire dataset. This fully automated process provided consistent, unbiased segmentation and quantification of astrocyte reactivity, reducing human error and expediting analysis. After segmentation, patches were reassembled to reconstruct full-size images for all 97 slides, with segmented astrocyte regions constrained within the predefined boundaries.

To reduce segmentation artifacts and false positives, morphological filtering was applied prior to morphometric analysis. Individual astrocytes were isolated from segmented GFAP-positive masks obtained from coronal pathology images. Custom MATLAB scripts were developed to automatically crop each astrocyte into individual image patches from the full coronal sections. Segmentations with high eccentricity, corresponding to elongated edge artifacts along tissue boundaries, were excluded to ensure that only true astrocytic structures were retained. In addition, MATLAB’s watershed segmentation algorithm, based on distance transforms, was applied to separate touching astrocytes that occasionally appeared merged in the 2D projections of segmented images, allowing individual cellular boundaries to be identified before cropping. To ensure accurate single-cell morphometry, segmentation artifacts were also removed using morphological filtering prior to SMorph skeletonization.

Next, atlas and pathology images were converted from pixels to micrometers using established pixel-to-µm conversion factors. For each animal, the MRI atlas was scaled to match post-perfusion brain dimensions by adjusting the anterior–posterior, medio–lateral, and dorso–ventral axes, accounting for inter-animal variability in brain size and processing-related shrinkage (Figure 3).

To facilitate precise overall and regional quantification of astrocyte reactivity, pathology slides were mapped onto a standard ferret brain MRI atlas [49] using a dedicated image registration pipeline. For each pathology slide, the ratio of the number of atlas slices to the number of pathology/tissue slices was calculated to determine a candidate set of atlas levels. A bounding box was fitted around the tissue region in each candidate MRI atlas slice, and the centroid of the box was calculated. A corresponding bounding box was drawn around the tissue boundaries in each pathology slide, as delineated during manual segmentation. Given the degree of tissue shrinkage introduced by chemical processing, each pathology bounding box was scaled by comparing its height and width to those of potential matching atlas slices within the candidate range. The centroid of each pathology box was then translated and overlaid onto the centroid of each atlas box in the candidate set. The atlas slice demonstrated the highest concordance in tissue boundaries and centroid alignment was selected via manual review as the corresponding anatomical level for each pathology slide. Once optimal registration was achieved, the coordinates of segmented astrocytes from each pathology image were transformed into standardized scaled atlas space, using the spatial relationship between each astrocyte segmentation cluster and the centroid of the appropriately scaled atlas and pathology slides. This process enabled the accurate assignment of reactive astrocytes and their morphological metrics to their respective anatomical regions, supporting a detailed analysis of regional reactivity and remodeling patterns in response to TBI.

The anatomical regions examined in this study were organized into major neuroanatomical regions and subregions based on the ferret brain MRI atlas[49]. Major neuroanatomical regions included white matter (WM), sulcus, gray matter (GM), cortical GM, and subcortical GM (Figure 3-C). It should be noted that in this atlas, the WM major region encompasses all white matter structures including corpus callosum, cingulum, fornix, cortex WM, cerebellum WM, and brainstem WM, among others. The atlas-defined GM major region specifically encompasses cerebellar GM, brainstem GM, and olfactory bulb, and is anatomically distinct from the cortical GM and subcortical GM categories, which are analyzed as separate major regions in this study. The sulcus category captures GFAP-positive astrocyte reactivity along the sulcal walls and depths of the gyrencephalic cortex, corresponding to the cortical tissue lining these infoldings rather than the sulcal space itself. Subregions were organized within their respective major region categories as follows: WM subregions included corpus callosum, cingulum, fornix, cortex WM, and cerebellum WM; GM subregions included cerebellum GM and brainstem; cortical GM subregions corresponded to individual gyral parcellations; and subcortical GM subregions included the amygdala, caudate, hippocampus, hypothalamus, putamen, and thalamus (Figure 3-D).

### Regional Analysis of Astrocyte Reactivity and Single-Cell Astrocyte Morphometric Analysis

The cropped astrocyte masks mentioned earlier were exported for morphometric analysis using the SMorph framework implemented in Python within a Jupyter Notebook environment. SMorph’s default built-in preprocessing and segmentation steps did not perform reliably for ferret astrocytes and were thus bypassed, and the pre-segmented masks generated by our in-house deep learning model were used directly as inputs in SMorph for skeletonization and feature extraction. Morphological features extracted by SMorph as presented in Figure 4, included the number and lengths of primary, secondary, tertiary, and quaternary branches, convex hull area, and astrocyte area. SMorph outputs were reported at the appropriate physical scale for quantitative analysis.

**Figure 4.**
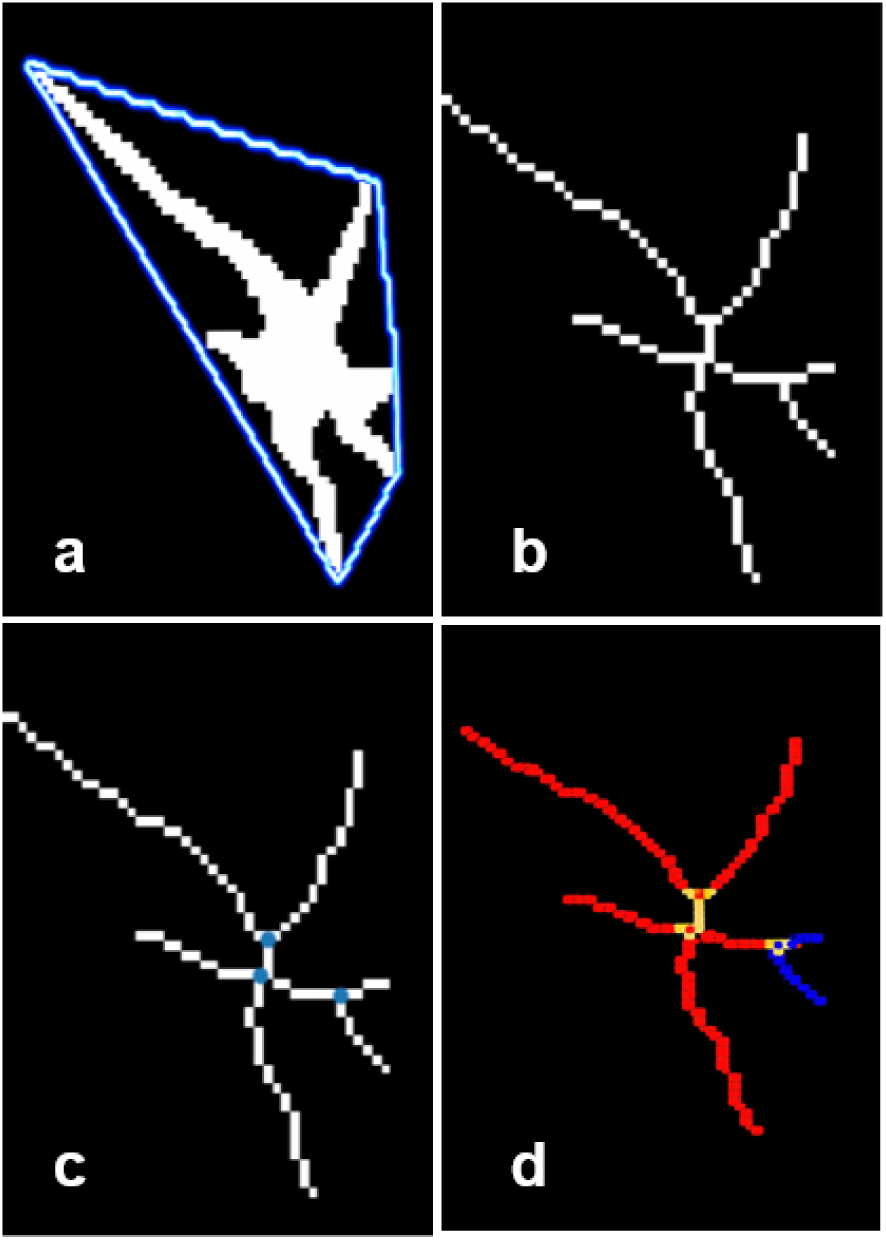
Astrocyte morphometric analysis using the SMorph pipeline: a) Convex hull b) Cell skeleton c) Branch detection d) Branch length

SMorph’s default built-in preprocessing and segmentation steps did not perform reliably for ferret astrocytes and were thus bypassed such that the our custom deep-learning model pre-segmented masks

All morphometric steps were executed using automated batch pipelines to ensure consistent processing across all samples. In total, 2,031,815 astrocytes from both sham and single severe TBI animals at 7 days post-injury were analyzed.

To quantify the proportional distribution of astrocyte reactivity within major neuroanatomical regions for each group (TBI or Sham) and determine whether the astrocyte reactivity distribution were similar or different between groups, the following formula was used:

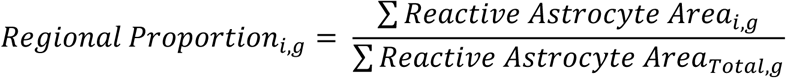

where *i* denotes major neuroanatomical region and *g* denotes group. This proportion represents the fraction of total astrocyte reactivity assigned to each region within a given experimental group, enabling comparative analysis of spatial distribution patterns between TBI and control cohorts.

Overall percentage of reactive astrocyte area for each animal was calculated as:

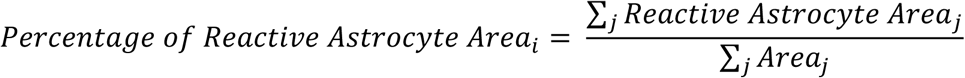

where *j* indexes pathology slices. This metric reflects the total area of reactive astrocytes across all slices, normalized by the total tissue area, thereby providing a whole-brain assessment of astrocyte reactivity independent of regional distribution.

Regional percentage of reactive astrocyte area within each anatomical area was computed as follows:

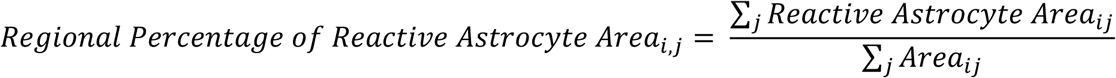

where *i* indexes anatomical regions, and *j* indexes slices. This calculation considers only atlas regions intersecting the mapped pathology images, thereby ensuring accurate regional quantification within the analyzed sample.

Mann-Whitney U tests were performed to compare overall and regional percentages of astrocyte reactivity between TBI and sham groups, with statistical significance defined as a p < 0.05, and p < 0.10 considered marginally significant.

Because morphometric data consisted of one observation per cell rather than a single summary value per animal, a separate statistical framework was employed to account for the hierarchical structure of the data. Morphometric metrics measured were analyzed at the single-cell level using linear mixed-effects models (LMM). For each parameter, experimental group was included as a fixed effect, and animal identity (ID) was modeled as a random intercept to account for the nested structure of astrocytes within animals. The model was defined as:

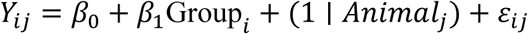

where *Y_ij_* represents the morphological metrics of astrocyte *i* from animal *j*.

Linear mixed-effects models were estimated using restricted maximum likelihood (REML), with denominator degrees of freedom approximated using the Satterthwaite method. This framework accounts for within-animal dependence and avoids pseudoreplication by treating the animal as the effective unit of inference. Intraclass correlation coefficients (ICC) were computed to quantify the proportion of variance attributable to between-animal differences. All analyses were performed in R, with statistical significance defined as *p* < 0.05, and *p* < 0.10 was considered marginally significant.

## Results

In the context of assessing the human relevance of the ferret brain for regional astrocyte analysis after TBI, our cross-species comparison showed that, in addition to previously reported morphological and functional similarities of astrocytes in ferrets and humans [89], the ferret brain displays regional fractional volumes (volume of each region divided by total brain volume) for major structures and TBI-relevant subregions that are much closer to human indices than rodent indices (Figure 1), indicating greater human–ferret than human–rodent similarity. For example, the volume fractions for WM, GM, cortical GM, subcortical GM, amygdala, brainstem, hippocampus, hypothalamus, thalamus, and corpus callosum are 21.5%, 18.0%, 42.3%, 13.9%, 0.2%, 5.6%, 3.6%, 1.2%, 3.5%, and 1.1%, respectively, in ferrets, closely matching human values (42.8%, 11.9%, 39.3%, 11.5%, 0.2%, 2.1%, 0.7%, 0.8%, 1.4%, 0.2%) and differing from mouse/rat ranges (9.1-10.5%, 20.9-22.7%, 27.9-36.1%, 26.7-28.8%, 2.2-3.0%, 6.3-9.8%, 5.9-7.4%, 2.5-2.8%, 4.2-4.7%, 1.5-3.6%) (Figure 1, Supplementary Table S1). These results demonstrate that the ferret, selected for this study, exhibits neuroanatomical characteristics and regional volume distributions that more closely resemble the human brain than traditional rodent models such as mice and rats, which are more commonly used in TBI research, while remaining less costly and requiring substantially less infrastructure than larger animal models such as pigs and primates.

The astrocyte segmentation AI model used in this study has been shown to accurately segment astrocyte reactivity across a range of imaging conditions encountered in GFAP-stained ferret brain tissue, including regions with normal, dense, or sparse astrocyte distributions, tissue border regions, and suboptimal imaging conditions such as blurring or dust artifacts [7]. As noted in the Methods section, pathology analysis was conducted on an average of 10 slides per brain, with corresponding locations in the ferret MRI atlas identified and pathology images overlaid on the atlas (Figure 5). These results demonstrate the mapping technique’s accuracy and effectiveness in aligning anatomical regions for regional analysis and confirm that this process was applied throughout the brain along the anteroposterior axis.

**Figure 5.**
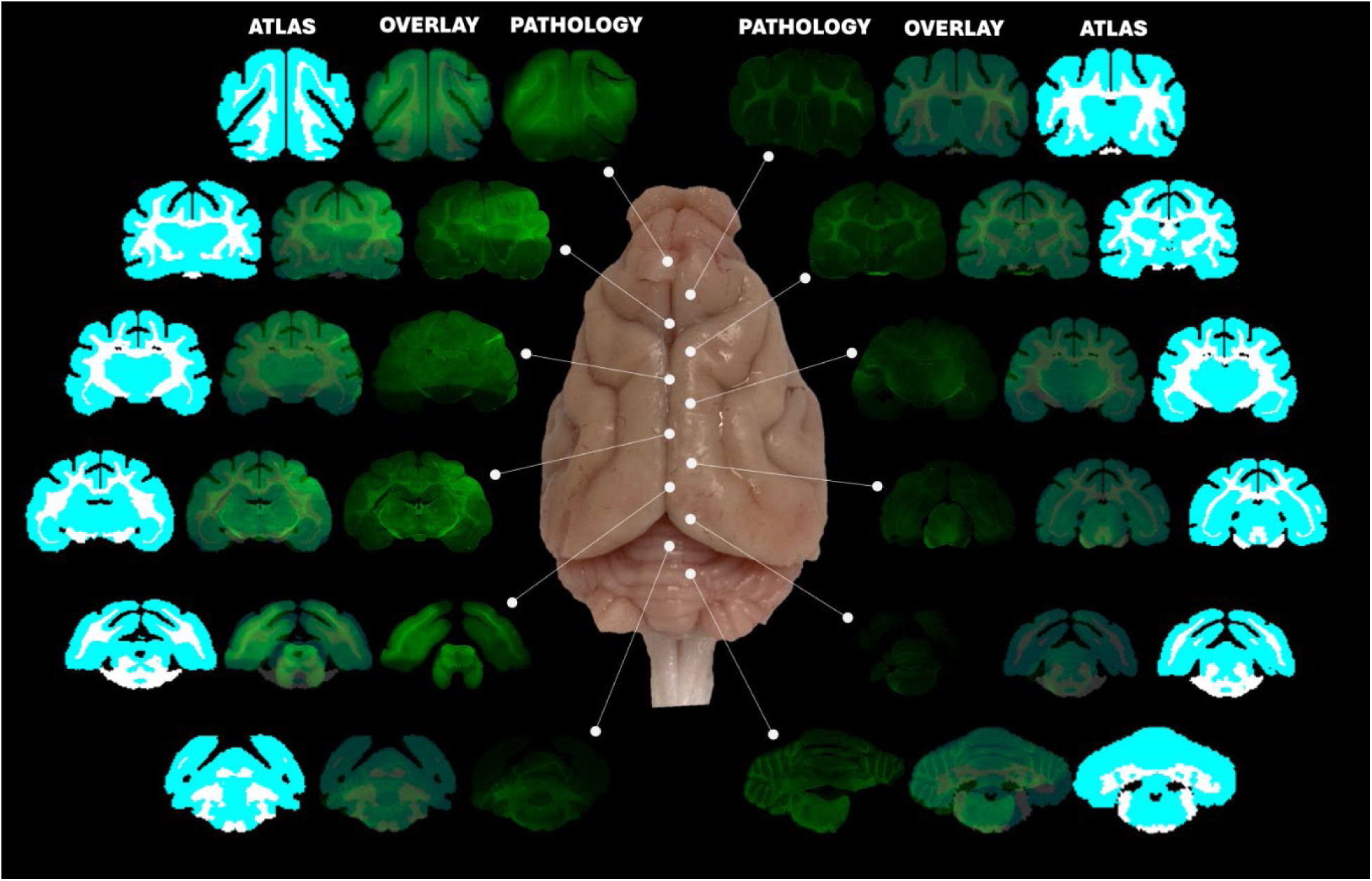
Example from a single ferret illustrating pathology slices aligned with their corresponding atlas slices, alongside atlas–pathology overlays. The consistent alignment across the anterior–posterior axis demonstrates accurate anatomical matching between pathology images and atlas slices.

Quantification of astrocyte reactivity across the entire brain revealed greater reactivity in the TBI group (average of 1.16%) compared with the sham group (0.95%) (Figure 6A), indicating a robust astroglial response elevated 7 days after TBI. Analysis of major neuroanatomical regions (Figure 6B) also showed that diffuse TBI elevates astrocyte reactivity across multiple regions, with particularly pronounced effects in GM (sham: 1.25% vs. TBI: 2.86%; p < 0.05). Analysis of 13 key brain subregions further revealed notable regional differences in astrocyte reactivity in the TBI and sham groups (Figure 6C). Cerebellum WM and brainstem showed the largest astrocyte reactivity differences, with TBI values of 2.82% and 1.82%, respectively, versus sham values of 0.96% and 0.58% (all p < 0.05). A marginal difference was observed in cerebellum GM (sham: 1.49% vs. TBI: 3.05%). Thalamus (sham: 1.14% vs. TBI: 1.59%), and Amygdala (sham: 0.41% vs. TBI: 1.04%) also showed notable astrocyte reactivity between sham and TBI groups, although these did not reach statistical significance. Additionally, corpus callosum, cortex WM, hippocampus, and hypothalamus exhibited higher reactivity in TBI than sham, although these differences were not statistically significant. A few regions showed slightly higher astrocyte reactivity in sham, most notably the putamen (sham: 3.36% vs. TBI: 1.85%). Figure 7 presents representative examples of astrocyte reactivity across key sub-regions for both groups, illustrating the differences between the TBI and sham groups across critical brain regions.

**Figure 6.**
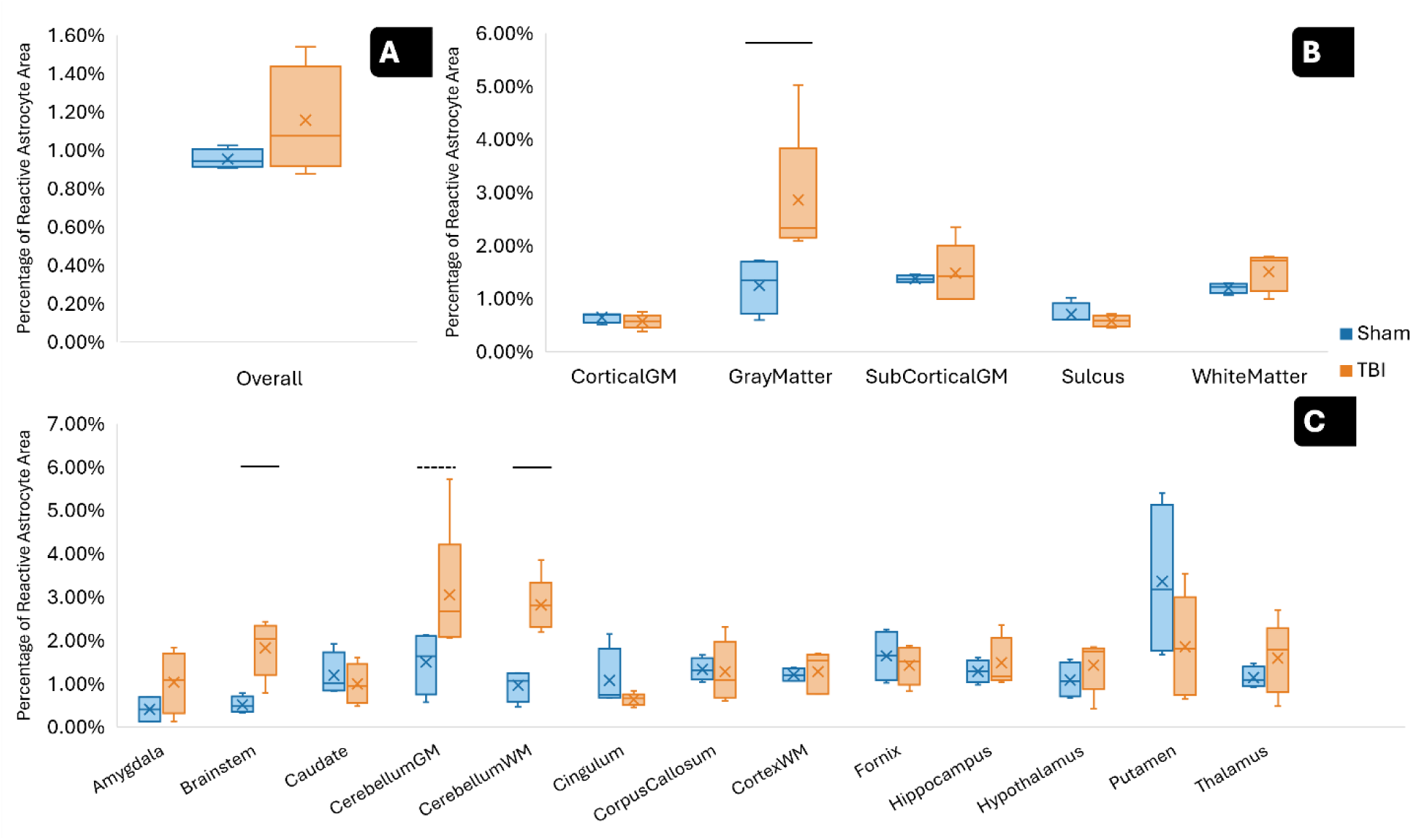
Comparison of percentage of astrocyte reactivity within each region between sham and TBI brains, shown for whole brain (A), major brain regions (B), and within key sub-regions (C). Significant (P < 0.05) and marginal significant (P < 0.1) differences between TBI and sham groups were shown with solid and dash lines, respectively. Overall, elevated astrocyte reactivity was observed in the TBI group compared to sham, with significant differences at the whole-brain level and in several regions and sub-regions.

**Figure 7.**
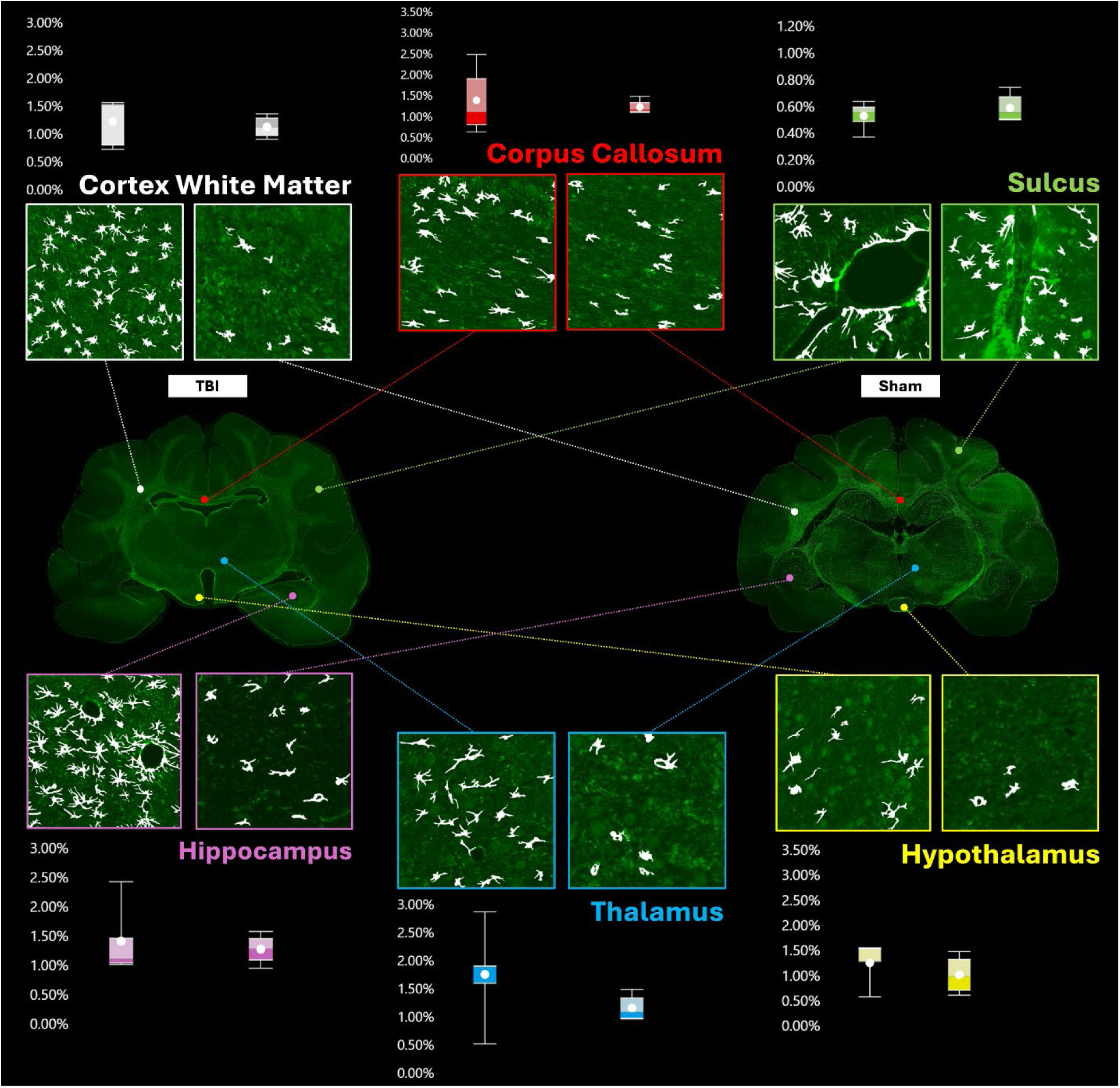
Representative sample visualizations and corresponding boxplots illustrating GFAP-positive astrocyte reactivity in key sub-regions for sham and TBI brains. Examples from cortex white matter, corpus callosum, sulcus, hippocampus, thalamus, and hypothalamus highlight region-specific activation patterns and increased astrocyte reactivity following TBI.

Figure 8 presents heatmaps of astrocyte reactivity across brain pathology slices from one TBI animal and one sham animal, each shown as a representative example of its respective group. The heatmaps illustrate the spatial distribution of reactive astrocyte area across 10 brain slices for both animal groups, clearly depicting areas with higher levels of astrocyte reactivity that correspond to localized astroglial responses following TBI. This visualization provides detailed insight into the spatial patterns of astrocyte reactivity resulting from diffuse TBI, observed across brain areas. By providing a comprehensive assessment of astrocyte reactivity in multiple slices per animal, the heatmaps enhance understanding of differential regional responses to diffuse TBI and support the findings from previous quantitative analyses reported in Figure 6.

**Figure 8.**
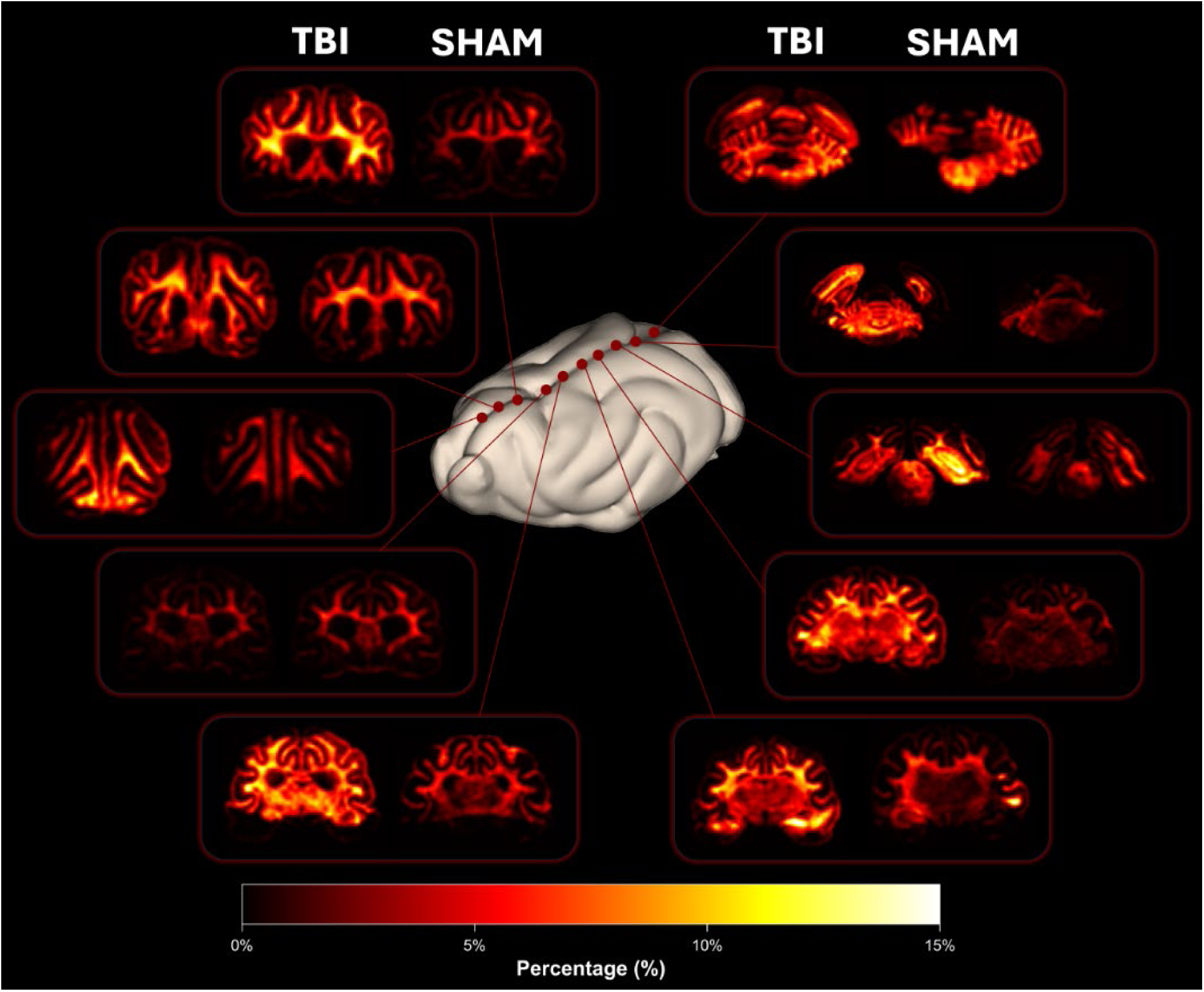
Heatmaps illustrate a sample of the spatial distribution of GFAP-positive astrocyte reactivity in TBI and sham groups. The maps demonstrate regional patterns of activation across 10 pathological slices, with the TBI animal exhibiting higher localized astrocyte reactivity.

Morphological analysis across overall, major regions, and subregions, summarized in Figure 9, revealed generally higher astrocyte morphological values, including cell area, convex hull area, number of forks, and number and length of branches, in the TBI group relative to the sham group. Branch length analysis demonstrated a consistent structural pattern in both groups, where primary branches exhibited the greatest length and progressively decreased toward quaternary branches. Among the analyzed features, LMM analysis revealed that, at the whole-brain level, convex hull area (sham: 386µm^2^ vs. TBI: 463µm^2^), astrocyte cell area (sham: 264µm^2^ vs. TBI: 293µm^2^), and secondary branches length (sham: 3.61µm vs. TBI: 3.97 µm) showed statistically significantly greater values in the TBI group, while tertiary (sham: 1.32µm vs. TBI: 1.48µm) and quaternary branch lengths (sham: 0.54µm vs. TBI: 0.61µm) were marginally greater in the TBI group compared with the sham group, consistent with overall astrocyte hypertrophy following TBI. Further analysis at main region and subregion levels revealed notable differences in gray matter, white matter, brainstem, cerebellum GM, cerebellum WM, and hippocampus. These regions exhibited the most pronounced morphological changes, showing significant or marginal differences across multiple parameters, including convex hull, astrocyte area, number of forks, and length of all types of branches. Even for parameters that did not reach statistical significance, astrocyte morphological values in these regions remained consistently higher in the TBI group compared to the sham group. All observed between-group differences are summarized in Figure 11. Also, ICC in LMM analysis were consistently low across all morphological parameters (ICC < 1.5%), indicating that most variability in astrocyte morphology arose from differences between individual cells rather than between animals.

**Figure 9.**
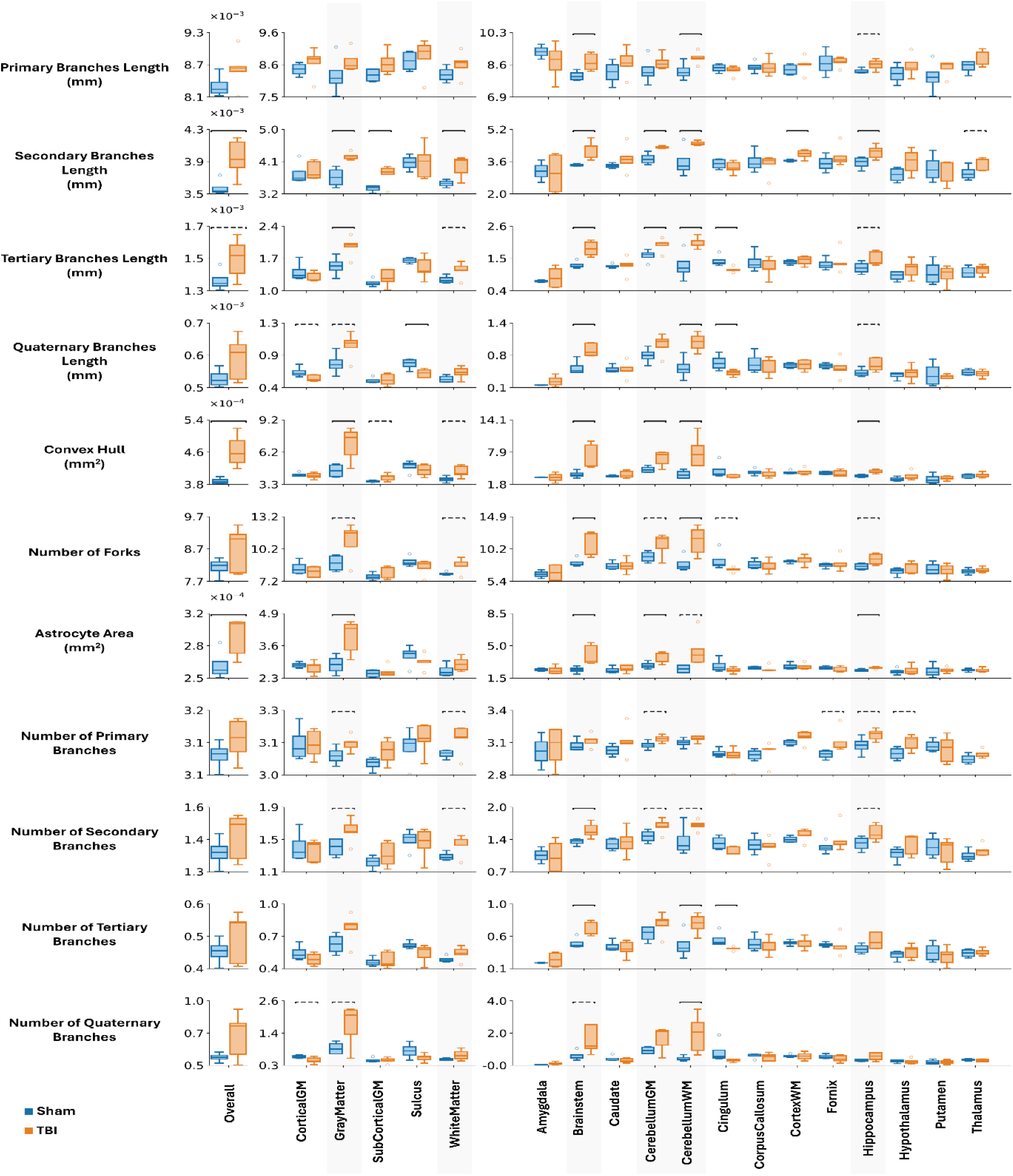
Astrocyte morphometric comparison between Sham (blue) and TBI (red) groups across overall, major regions, and subregions. Boxplots show distributions of astrocyte morphological parameters including branch lengths, convex hull area, number of forks, and branch counts across hierarchical orders. Blue represents TBI and orange represents Sham. Black asterisks indicate statistically significant differences (P < 0.05), and gray asterisks indicate marginal significance (P < 0.1).

## Discussion

### Ferrets provide a human-like model for investigating whole-brain and region-specific astrocyte pathology after diffuse TBI

Rodent models (mice and rats) and, to a lesser extent, pigs are the most widely used experimental systems for TBI because rodents are accessible, cost-effective, and supported by extensive methodological infrastructure, whereas pigs offer closer anatomical similarity to the human brain. However, the lissencephalic structure and relatively low white-matter content of rodent brains differs substantially from the gyrencephalic, white-matter–rich human brain, limiting their ability to recapitulate diffuse injury processes such as axonal damage in long-range tracts [3, 34]. In contrast, large gyrencephalic species like pigs reproduce human neuroanatomy and TBI biomechanics but require substantial resources, specialized housing, and intensive veterinary support, and typically permit only modest sample sizes, limiting their practicality for systematic analyses. Together with our reginal pathological findings, previously reported human-ferret similarity in astrocyte morphology and function, and the anatomical comparisons in Figure 1 and Supplementary Table 1, these critical attributes support the ferret as an intermediate model that combines key neuroanatomical and astrocyte features of the human brain with the practicality of a small mammalian model organism. Ferrets possess a gyrencephalic cortex with prominent gyri and sulci, a ventrally positioned hippocampus, and regional fractional volumes that closely parallel human values, than those of mice and rats, including higher WM and cortical GM fractions and relatively lower total GM and subcortical GM fractions. In addition, several TBI-relevant subregions, such as the amygdala, hippocampus, hypothalamus, and corpus callosum, have volume fractions that are smaller and therefore more comparable to humans than to rodents, suggesting that the distribution of tissue and cellular deformation during TBI, and the relative susceptibility of these structures to injury, can be modeled more closely to the human condition in ferrets than rodents. The gyrencephalic architecture and a more human-like WM organization are particularly important for diffuse, rotation-driven TBI, where strain concentrates in deep tracts and GM–WM interfaces [42, 43, 45], and drives regionally heterogeneous TBI pathology [101], such as astrocyte reactivity patterns, similar to those observed in human patients. Collectively, anatomical and astrocytic similarities, combined with lower cost and reduced infrastructure requirements compared to large-animal models, position the ferret as a valuable platform for studying diffuse TBI. The combination of translational relevance and practical feasibility makes the ferret particularly well suited for whole-brain, region-specific quantification of astrocyte pathology, as implemented in this study.

### GFAP-positive astrocyte reactivity provides a spatially resolved measure of diffuse TBI progression

Beyond animal model selection, the choice of pathological markers strongly influences how TBI progression is captured. Astrocytes are among the earliest and most robust responders to brain injury, undergoing rapid hypertrophic remodeling and transcriptional reprogramming and engaging in signaling with neurons, microglia, and the cerebrovascular compartment as part of both primary and secondary injury cascades[14, 15, 20, 57, 73, 77, 116]. In contrast to pathological readouts that primarily index acute and primary injury such as traumatic axonal injury, which often peak within the first 24 hours after impact and then decline, astrocyte pathology evolves over a longer time course and is observed across acute, subacute, and chronic phases of TBI [54, 112]. Reviews of TBI time courses emphasize that many axonal injury markers are most prominent early, whereas astrocyte reactivity and glial scar formation persist and can be detected weeks to months and even observed a year after injury [2, 103, 105]. Several studies further indicate that GFAP-positive astrocyte reactivity frequently reaches a subacute peak around 7 days post-injury, providing a window for capturing both established pathology and emerging remodeling processes [14, 15, 31, 50, 65, 66, 85, 104, 113, 118]. In contrast to the ferret CHIMERA TBI study by Krieg et al. [62], which examined astrocyte reactivity at 24 hours post-injury in a toddler age group of ferrets, the present study assessed astrocyte pathology in adult-age ferrets at this 7-day subacute peak to characterize injury progression beyond the immediate acute phase and to capture the onset of longer-term pathological and repair processes [70, 118].

Additionally, astrocyte responses to injury are increasingly recognized as spatially heterogeneous and strongly influenced by the underlying injury mechanism, with diffuse TBI producing widespread biomechanical strain and region-dependent astrocyte responses rather than the localized peri-lesional astrocytic gradients typically observed in focal injury models [80]. Within this framework, reactive astrocytes have context-dependent effects: acutely they may help contain tissue damage, stabilize the blood–brain barrier, buffer excitotoxic transmitters, and support repair processes and homeostatic functions, whereas sustained or excessive activation is associated with glial scar formation, chronic neuroinflammation, and reduced axonal plasticity [13, 15, 82, 103–105]. Given this prolonged, region-specific, and functionally diverse response, GFAP-based astrocyte reactivity provides a temporally durable and spatially informative readout of sustained astroglial engagement at the subacute peak of gliosis, making it well suited for whole-brain, region-specific quantification implemented in this study. Nevertheless, the stability of GFAP immunostaining across injury phases, while advantageous for detecting persistent reactivity, does not by itself resolve the dynamic molecular state transitions of astrocytes that occur across the TBI timeline[2, 80].

GFAP, an intermediate filament protein highly enriched in mature astrocytes, was selected as the primary histological marker in this study because it is one of the most extensively characterized indicators of reactive astrogliosis in experimental and clinical TBI. GFAP is robustly upregulated following central nervous system (CNS) injury and shows relatively sustained expression during the subacute post-injury period analyzed here, while also providing reliable and technically straightforward immunostaining in paraffin-embedded brain tissue. Importantly, GFAP directly reflects the cytoskeletal remodeling associated with reactive astrocyte hypertrophy and provides clear visualization of astrocytic morphology, including process extension and structural reorganization[2, 5, 24, 115]. Compared with other astrocyte markers such as S100β and ALDH1L1, which are more suitable for identifying astrocyte cell bodies or total astrocyte populations, GFAP is particularly well suited for assessing injury-induced astrocyte reactivity[33, 55, 105]. Likewise, compared with vimentin, nestin, and other injury-associated astrocyte markers that are less specific for mature reactive astrocytes, GFAP has been more broadly validated in TBI models and remains the most widely used marker for reactive astrogliosis [28, 55, 85, 105]. Therefore, GFAP was selected as the histological marker of astrocyte reactivity in the present study because it remains the most direct and well-established indicator of reactive astrogliosis following TBI. In this context, regionally resolved GFAP-positive astrocyte pathology provides an integrative and spatially resolved measure of how diffuse biomechanical loading translates into local vascular, metabolic, and synaptic disturbances across distinct brain regions, thereby offering an informative pathological readout for tracking diffuse TBI progression and establishing a baseline for evaluating future therapeutic interventions, including astrocyte-targeted strategies.

### Deep-learning-based astrocyte segmentation and morphological analysis enable scalable astrocyte pathology assessment

Adopting an AI-driven model for automatic astrocyte reactivity segmentation strengthens this approach by enabling whole-brain analysis at a scale not achievable through manual or threshold-based methods, which are time-consuming, labor-intensive, and/or prone to variability [4, 19, 29, 47, 71, 93, 103, 119]. Deep-learning-based methods have consistently demonstrated superior segmentation performance on biomedical images, including astrocytes [59], offering greater accuracy, consistency, and efficiency. In our previous work, we identified the U-Net++ architecture paired with a VGG19 backbone as the best-performing configuration for segmentation of GFAP-stained astrocyte reactivity in fluorescence microscopy images [7]. Integrating this model with atlas-guided regional mapping enabled quantitative, high-fidelity analysis across the full coronal extent of the brain and across anatomically defined subregions, yielding a more comprehensive whole-brain and regional assessment of astrocyte reactivity than traditional approaches that are qualitative or, when quantitative, restrict analysis to a limited number of slices and/or regions and may fail to capture distributed, network-level patterns of astrocyte reactivity and remodeling across brain regions [25, 56, 94].

### Whole-brain and region-/subregion-specific astrocyte reactivity following diffuse TBI

Applying this framework across coronal slices from the brainstem to the frontal lobe along the anteroposterior axis revealed greater astrocyte reactivity in the TBI group than in the sham group across the entire brain, but nonuniformly distributed across different regions (Figures 6-8). This pattern is consistent with astrocyte reactivity as a core glial response to TBI [14, 15, 76, 105], but also indicates that diffuse, rotation-driven injury does not produce a uniform, brain-wide increase in GFAP signal. Diffuse TBI is characterized by rapid head rotation that induces deformation of brain tissue, axons, and supporting structures across multiple regions, resulting in widespread pathology and disruption of neural networks, in contrast to focal TBI, which primarily causes localized injury [40, 42, 102].

Among the five major regions studied, the GM region showed a statistically significant increase in GFAP-positive reactivity, with approximately a threefold higher percentage of reactive astrocyte area in the TBI group than in the sham group, whereas the WM region exhibited an almost two-fold higher mean value in the TBI group compared to the sham group that did not reach statistical significance. A portion of this regional distribution is likely driven by the widespread, rotation-induced mechanical loading characteristic of diffuse TBI, which imposes inertial deformation/strain on both long-range white-matter tracts and GM–WM interfaces intersected by major fiber pathways. Mechanical loading, however, is only the initiating event and does not fully explain the pattern of astrocyte reactivity observed at 7 days post-injury; secondary injury cascades further shape these responses over time. Disruption of the blood–brain barrier (BBB) can permit extravasation of blood-derived proteins such as fibrinogen into the parenchyma, triggering astrocyte activation in perivascular domains and contributing to partial restoration of barrier integrity [18, 23, 99], while excitotoxicity, ischemia, and oxidative stress further contribute to ongoing tissue damage and astrocyte reactivity [9, 108].

Within this framework, the greater and statistically robust increase in astrocyte reactivity in GM likely reflects the combined effects of GM–WM interface strain, high synaptic density, and the predominance of protoplasmic astrocytes in GM, which are intimately associated with neuronal cell bodies, synapses, and capillaries and may respond strongly to synaptic and metabolic disturbances [8, 32, 60, 69, 72, 77, 113]. On the other hand, WM is dominated by fibrous astrocytes with less branched processes along myelinated axon bundles, in a microenvironment where axonal stretch, myelin disruption, and metabolic stress are prominent after diffuse TBI. These features likely make WM astrocytes highly responsive to axonal and myelin injury; accordingly, GFAP-positive area was greater (approximately two-fold) in WM in the TBI group compared to sham, although this difference did not reach statistical significance. In subcortical gray matter, which is embedded within or adjacent to major fiber bundles, we observed slightly larger astrocyte reactivity values in TBI than sham, but with region-dependent variability, suggesting that local connectivity and tract architecture may further modulate the relationship between diffuse loading and astrocyte activation. Like subcortical-GM, GFAP-positive astrocyte reactivity in sulcus and cortical GM regions was very similar between TBI and sham, which may be partly explained by regional differences in effective strain or injury loading and partly by distinct astrocyte response phenotypes. In mild diffuse TBI, atypical astrocytes have been described in BBB-disrupted microenvironments, where they rapidly lose key homeostatic proteins and astrocyte–astrocyte coupling without showing robust increases in classical reactive markers such as GFAP, such that substantial astrocyte pathology may not necessarily present as a larger GFAP-positive area and astrocyte reactivity [37, 97]. Because BBB disruption after TBI is known to be spatially heterogeneous and we did not quantify BBB integrity or non-GFAP astrocyte markers in this study, a potential contribution of such atypical responses to the lack of detectable GFAP differences in sulcus and cortical GM and subcortical GM is a plausible but unproven mechanism that requires future investigation.

Sub-regional analyses also demonstrated substantially greater astrocyte reactivity across several key sub-regions in response to TBI compared with sham, particularly in the brainstem and cerebellum WM and GM, which showed significant or marginal group differences, whereas thalamus, hypothalamus, amygdala, and cortex WM exhibited only modest increases in TBI relative to sham. These patterns suggest region-specific astroglial responses to diffuse injury and align with prior clinical and preclinical evidence of widespread network disruption following TBI. The brainstem is a primary site of axonal injury in diffuse TBI because its long descending and ascending fiber tracts are highly susceptible to large deformation/strain during rapid head rotation. Astrocytes in the brainstem play a critical role in supporting the function of long-range axonal circuits and maintaining the local microenvironment of neurons responsible for vital autonomic and homeostatic functions; their activation and structural remodeling therefore likely reflect both direct mechanical perturbation and secondary inflammatory signaling arising from widespread axonal damage in this region [29, 102]. In cerebellum GM, regions enriched in Purkinje neurons have high synaptic density and metabolic demand, making them particularly vulnerable to synaptic disruption and excitotoxic injury; astrocyte remodeling in this sub-region likely reflects attempts to restore neurotransmitter balance and maintain local circuit activity. By comparison, cerebellum WM contains densely packed myelinated axonal tracts that are highly vulnerable to axonal strain in diffuse TBI, and astrocyte alterations in this region are consistent with responses to axonal injury, ionic imbalance, and inflammation within fiber-rich pathways [22, 36, 62, 83, 105].

This sub-regional pattern is at least partly explainable by the distribution of mechanical loading and tissue deformation in our sagittal rotational diffuse TBI model. Head rotation about the neck in the sagittal plane is expected to generate relatively large tissue and axonal longitudinal strain along the cerebellar peduncular tracts connecting the brainstem and cerebellum, while more centrally located structures such as the thalamus, hypothalamus, and amygdala likely experience lower deformation magnitudes, consistent with the more modest TBI-versus-sham astrocyte reactivity differences observed in these regions. Structures such as the putamen, caudate, cingulum, and corpus callosum, which are more centrally located and lie closer to the mechanical plane of sagittal rotation, may undergo comparatively less deformation in this loading configuration, or may experience dominant deformation components that are not aligned with axonal fibers (for example, within the corpus callosum). These features could contribute to the small or absent increases in GFAP-positive astrocyte area, and in some subregions, TBI values that are similar to or even lower than sham values observed in our dataset. By contrast, axial and coronal head rotations are expected to shift peak tissue and tract-oriented strains toward more central and deep structures, including midline white-matter pathways, implying that different rotational axes may preferentially engage distinct subcortical astrocyte populations and produce larger longitudinal deformation in regions such as the corpus callosum. Although a systematic evaluation of this biomechanical explanation will require detailed finite-element reconstruction of the present ferret CHIMERA experiments, and efforts are underway in our laboratory to perform such simulations, our previous computational work in large-animal rotational TBI models has shown that traumatic axonal injury locations colocalize with regions of elevated tissue and tract-oriented strain [42, 43, 45], indirectly supporting the proposed link between regional deformation and astrocyte reactivity. In addition to these biomechanical considerations, astrocyte responses are highly phenotype- and context-dependent, so regional differences in the prevalence of atypical or GFAP-low astrocyte states, as well as in BBB disruption, vascular architecture, baseline astrocyte density, and the timing of astrocyte activation at 7 days post-injury, are also likely to contribute to the magnitude and variability of GFAP changes observed across subregions.

### Whole-brain and region-/subregion-specific astrocyte morphological changes following diffuse TBI

While GFAP upregulation is widely used as a hallmark of reactive astrogliosis, astrocyte responses to injury are increasingly recognized as heterogeneous and context-dependent, particularly with respect to injury type and the local tissue environment [8, 32, 80]. Previous work suggests that astrocytes may adopt multiple distinct reactive states following injury, and that associated morphological phenotypes can be characterized in greater detail when bulk GFAP expression is complemented by quantitative morphometric analysis [33, 80], highlighting the value of morphometry as an additional readout alongside traditional marker-based approaches. Our quantitative morphometric analysis (Figure. 10) revealed several significant structural changes following injury: astrocytes in the TBI group exhibited increased convex hull area, astrocyte cell area, and distal process length compared with the sham group, at the whole-brain level, as well as in several major regions and subregions, indicating expansion of the spatial domain occupied by individual astrocytes. Convex hull area represents the outer boundary of the astrocytic arbor and serves as a proxy for the effective spatial domain occupied by the cell; increases in this parameter are typically interpreted as hypertrophic remodeling or territorial expansion [8, 27, 98]. Increases in astrocyte area further corroborate that individual astrocytes became larger following TBI, consistent with the hallmark morphological feature of reactive astrogliosis, namely cellular hypertrophy accompanied by swelling of the cell body and thickening of processes [8, 14, 105]. Such expansion of astrocyte territory and elongation of intermediate-order processes is expected to alter the extent and pattern of astrocytic coverage of synapses and microvessels, with potential consequences for synaptic regulation, neurovascular coupling, and local excitatory–inhibitory balance [23, 78].

In addition, astrocytes displayed elongation of proximal and distal processes. Specifically, secondary branch length was the most consistently and significantly elevated morphometric parameter following TBI, both at the whole-brain level and across multiple regions and subregions. Secondary processes elongate to expand the surveillance territory of each astrocyte and increase coverage of synapses and vasculature over a wider domain [114]. This interpretation is supported by the concurrent significant increase in convex hull area and astrocyte cell area, which together indicate that the overall spatial domain of astrocytes expanded, and that this expansion appears to be driven more by elongation of intermediate-order processes rather than by additional branching of the most distal processes.

Across regions where significant or marginal branch differences were detected, changes in branch length parameters were more consistent and reached statistical significance more frequently than changes in branch count parameters at equivalent hierarchical levels, indicating that astrocytes respond more through hypertrophy and elongation of existing processes rather than through a large increase in branching complexity [106, 114], although increases in branch number at different hierarchical levels were also observed to a lesser degree. This elongation-dominant response is characteristic of the mild-to-moderate hypertrophic form of reactive astrogliosis that occurs in diffuse mild TBI, in which astrocytes remain more within their individual territorial domains and extend their processes to compensate for tissue perturbation, while exhibiting modest increases in branching and proliferation compared with the more pronounced proliferative responses typically reported in severe or focal injuries [10, 106].

Consistent with the GFAP-based regional findings (Figure. 6), the regions and subregions showing the most notable astrocyte reactivity also demonstrated the most extensive morphological remodeling, across all or most of morphometric parameters studied, including convex hull, astrocyte areas, branch length and number at different hierarchical levels. GM exhibited the most statistically consistent pattern of morphological change among all major regions examined, with significant differences in secondary branch length, convex hull area, and astrocyte cell area, as well as marginal differences in tertiary branch length, number of secondary branches, number of forks, quaternary branch length, number of quaternary branches, and number of primary branches. This concentration of morphological differences in GM is consistent with the mechanical and neurochemical vulnerabilities of this region described earlier. WM also exhibited a similar pattern for most morphological metrics. At the subregion level, the brainstem, cerebellum GM, cerebellum WM, and hippocampus demonstrated the most pronounced morphological differences. The astrocyte morphological remodeling in the brainstem and cerebellum GM and WM in the TBI group relative to the sham group is aligned with their GFAP-positive astrocyte reactivity responses described earlier and is consistent with their susceptibility to axonal injury, ionic imbalance, and circuit disruption in diffuse TBI [10, 29, 30, 39, 102, 105]. In these three subregions, the combination of increased convex hull area, astrocyte area, and branch elongation suggests that astrocytes expand their territorial coverage along vulnerable long-range fiber tracts and synapse-dense cerebellar layers, consistent with heightened demands for ion buffering, neurotransmitter clearance, and metabolic support. This structural hypertrophy provides a cellular correlate for the strong GFAP-positive astrocyte reactivity in these regions and supports the idea that astrocytes help stabilize long-range pathways subjected to high strain under sagittal rotational loading.

Although the hippocampus did not show a statistically significant increase in GFAP-positive astrocyte reactivity, it was among the most extensively remodeled subregions in the morphometric analysis, with significant differences in convex hull area and astrocyte area and marginal differences across several additional parameters. This region is recognized as one of the most injury-sensitive brain structures following TBI, owing to its exceptionally high synaptic density, dense expression of glutamate receptors, and critical role in memory consolidation and spatial navigation [9, 39]. The presence of marked morphological changes in the absence of a corresponding significant increase in GFAP-positive area indicates that structural remodeling of astrocyte processes represents a partially independent dimension of the astroglial injury response that is not fully captured by bulk GFAP measurements. These results reinforce the value of combining molecular markers with quantitative morphological analysis to capture the full spectrum of astrocyte responses following injury. In the hippocampus, such remodeling may be particularly relevant because changes in astrocyte territory and process architecture are expected to influence synaptic integration, plasticity, and neurovascular coupling within circuits that underlie TBI-related memory and spatial navigation deficits.

The low intraclass correlation coefficients observed across all morphometric parameters (ICC < 1.5%) confirm that astrocyte morphological heterogeneity is predominantly a cell-level phenomenon, and that animals within the same experimental group behaved consistently with one another, supporting the reliability of the group-level morphometric comparisons.

### Limitations

The present work implements an AI-based framework for astrocyte segmentation, morphometry, and atlas-guided regional mapping, but several limitations should be noted. First, the analysis is restricted to a single subacute time point; future studies incorporating acute and chronic intervals will be needed to obtain a temporally resolved picture of astrocyte reactivity and its relationship to behavioral outcomes. Histological assessment was also limited to GFAP immunostaining, without additional markers of TBI pathology such as β-APP for axonal injury, Iba-1 for microglial activation, or fibrinogen for BBB disruption, and reliance on GFAP alone does not distinguish protective from maladaptive astrocytic states. Future work could expand this framework by incorporating additional astrocyte markers (e.g., vimentin or A1/A2-associated markers [30, 61]) and by extending quantitative mapping to other TBI-relevant pathologies such as axonal injury (e.g., β-APP or neurofilament staining [42]) and BBB disruption (e.g., fibrinogen or IgG leakage [11, 92]). The present study was conducted exclusively in male ferrets, so sex as a biological variable was not examined; future studies including female animals will be necessary to determine whether the regional astrocyte response patterns identified here are sex-dependent. In parallel, detailed finite element simulations of the ferret CHIMERA configuration could provide quantitative maps of regional tissue and tract-oriented strain to rigorously test the proposed links between mechanical loading and tissue deformation patterns and astrocyte responses, and applying the same atlas-registered approach to additional markers of axonal injury, BBB disruption, and diverse astrocyte states would enable a more comprehensive characterization of TBI pathology over time.

## Conclusion

Taken together, this work establishes the ferret CHIMERA model, combined with AI-based segmentation and atlas-guided regional mapping, as a practical and human-relevant platform for whole-brain quantification of astrocyte pathology in diffuse, rotation-driven TBI. Beyond the overall increase in astrocyte reactivity and morphological remodeling at the whole-brain level, gray matter overall and specific subregions, particularly the brainstem and cerebellar WM and GM, showed the most pronounced and statistically significant increases in GFAP-positive astrocyte reactivity and hypertrophic morphological remodeling following rapid sagittal rotation, with white matter showing a qualitatively parallel but less marked trend. This spatial pattern is at least partly explainable by expected relatively large tissue and axonal strains at GM–WM interfaces and along long-range brainstem–cerebellar fiber tracts in this sagittally rotated TBI model, underscoring the key role of biomechanical loading, tissue deformation, and axonal longitudinal strain in shaping the pathological consequences of TBI, even though such strain patterns remain to be quantified directly in future finite element studies. Together with the high synaptic and metabolic demand of GM and the differing predominance of protoplasmic versus fibrous astrocytes, which differentially respond to synaptic, axonal, and vascular disturbances, these factors likely contribute to the observed regional profile of astrocyte activation. In contrast, several centrally located subcortical structures and the hippocampus display more modest or heterogeneous changes in GFAP-positive area, even as the hippocampus still shows substantial astrocyte hypertrophy, underscoring that structural remodeling represents a partially independent dimension of the astroglial response. By integrating regional GFAP mapping with large-scale morphometry, this framework suggests relationships between expected mechanical loading and tissue deformation patterns and spatially specific astrocyte phenotypes and underscores the need for future multimodal and computational studies to characterize, and ultimately inform interventions for, region-specific vulnerability and repair following diffuse TBI.

## Ethics Approval

All animal procedures were approved by the Institutional Animal Care and Use Committee (IACUC) of the University of Texas at San Antonio (protocol #MF001) and conducted in accordance with the National Institutes of Health guidelines for the care and use of laboratory animals. Anesthesia was maintained using isoflurane, and euthanasia was performed using a pentobarbital-based solution administered via intracardiac injection, consistent with the American Veterinary Medical Association (AVMA) Guidelines for the Euthanasia of Animals (2020). No client-owned animals were used in this study. Consent to participate is not applicable.

## Consent for publication

Not applicable

## Availability of data and material

The AI model used in this study is on our lab GitHub, https://github.com/memar-lab/AutomaticAstrocyteSegmentation. The datasets generated and analyzed during this study are available from the corresponding author on request.

## Acknowledgments

Research reported in this publication was supported by National Science Foundation (CAREER # 2541217), start-up funding from the principal investigator, Memar, as well as funding from the Connecting through Research Partnerships (Connect) program sponsored by UT San Antonio. The authors also acknowledge the computational support provided by the High-Performance Computing Center (ARC) at the University of Texas at San Antonio. The views expressed are solely those of the authors and do not represent those of any funding sources or their affiliates.

## References

1 Abstracts Neurotrauma 2024 San Francisco, California. Journal of Neurotrauma 0: A-1-A-121 Doi 10.1089/neu.2024.41112.abstracts

2 Abou-El-Hassan H, Yahya T, Zusman BE, Albastaki O, Imkamp HT, Ye JJ, Percopo F, Christenson JR, Al Mansi MH, Lu K-J (2025) Astrocyte activation persists one year after TBI: a dynamic shift from inflammation to neurodegeneration. Communications Biology 8: 1745

3 Ackermans NL, Varghese M, Wicinski B, Torres J, De Gasperi R, Pryor D, Elder GA, Gama Sosa MA, Reidenberg JS, Williams TM (2021) Unconventional animal models for traumatic brain injury and chronic traumatic encephalopathy. Journal of neuroscience research 99: 2463–2477

4 Adelson PD, Jenkins LW, Hamilton RL, Robichaud P, Tran MP, Kochanek PM (2001) Histopathologic response of the immature rat to diffuse traumatic brain injury. Journal of neurotrauma 18: 967–976

5 Amlerova Z, Chmelova M, Anderova M, Vargova L (2024) Reactive gliosis in traumatic brain injury: a comprehensive review. Frontiers in Cellular Neuroscience 18: 1335849

6 Bagherian A, Abbasi Ghiri A, Ramzanpour M, Wallace J, Elashy S, Seidi M, Memar M (2025) Position-based assessment of head impact frequency, severity, type, and location in high school American football. Frontiers in Bioengineering and Biotechnology 12: 1500786

7 Bagherian A, Rahman Aranya OR, Kosub A, Redington M, Desai K, Memar M (2025) Comprehensive benchmarking of deep learning approaches for automated astrocyte segmentation in traumatic brain injury. Journal of Neuropathology & Experimental Neurology: nlaf114

8 Baldwin KT, Murai KK, Khakh BS (2024) Astrocyte morphology. Trends in cell biology 34: 547–565

9 Baracaldo-Santamaría D, Ariza-Salamanca DF, Corrales-Hernández MG, Pachón-Londoño MJ, Hernandez-Duarte I, Calderon-Ospina C-A (2022) Revisiting excitotoxicity in traumatic brain injury: from bench to bedside. Pharmaceutics 14: 152

10 Bardehle S, Krüger M, Buggenthin F, Schwausch J, Ninkovic J, Clevers H, Snippert HJ, Theis FJ, Meyer-Luehmann M, Bechmann I (2013) Live imaging of astrocyte responses to acute injury reveals selective juxtavascular proliferation. Nature neuroscience 16: 580–586

11 Bharadwaj VN, Rowe RK, Harrison J, Wu C, Anderson TR, Lifshitz J, Adelson PD, Kodibagkar VD, Stabenfeldt SE (2018) Blood–brainbarrier disruption dictates nanoparticle accumulation following experimental brain injury. Nanomedicine: Nanotechnology, Biology and Medicine 14: 2155–2166

12 Breedlove EL, Robinson M, Talavage TM, Morigaki KE, Yoruk U, O’Keefe K, King J, Leverenz LJ, Gilger JW, Nauman EA (2012) Biomechanical correlates of symptomatic and asymptomatic neurophysiological impairment in high school football. Journal of biomechanics 45: 1265–1272

13 Burda JE, Bernstein AM, Sofroniew MV (2016) Astrocyte roles in traumatic brain injury. Exp Neurol 275: 305–315

14 Burda JE, Bernstein AM, Sofroniew MV (2016) Astrocyte roles in traumatic brain injury. Experimental neurology 275: 305–315

15 Burda JE, Sofroniew MV (2014) Reactive gliosis and the multicellular response to CNS damage and disease. Neuron 81: 229–248

16 Calabrese E, Badea A, Coe CL, Lubach GR, Shi Y, Styner MA, Johnson GA (2015) A diffusion tensor MRI atlas of the postmortem rhesus macaque brain. Neuroimage 117: 408–416

17 Capizzi A, Woo J, Verduzco-Gutierrez M (2020) Traumatic brain injury: an overview of epidemiology, pathophysiology, and medical management. Medical Clinics 104: 213–238

18 Cash A, Theus MH (2020) Mechanisms of blood–brain barrier dysfunction in traumatic brain injury. International journal of molecular sciences 21: 3344

19 Cekanaviciute E, Buckwalter MS (2016) Astrocytes: integrative regulators of neuroinflammation in stroke and other neurological diseases. Neurotherapeutics 13: 685–701

20 Chen C, Zhong X, Smith DK, Tai W, Yang J, Zou Y, Wang L-L, Sun J, Qin S, Zhang C-L (2019) Astrocyte-specific deletion of Sox2 promotes functional recovery after traumatic brain injury. Cerebral Cortex 29: 54–69

21 Chen Z, Shin D, Chen S, Mikhail K, Hadass O, Tomlison BN, Korkin D, Shyu C-R, Cui J, Anthony DC (2014) Histological quantitation of brain injury using whole slide imaging: a pilot validation study in mice. PLoS One 9: e92133

22 Cheng WH, Martens KM, Bashir A, Cheung H, Stukas S, Gibbs E, Namjoshi DR, Button EB, Wilkinson A, Barron CJ (2019) CHIMERA repetitive mild traumatic brain injury induces chronic behavioural and neuropathological phenotypes in wild-type and APP/PS1 mice. Alzheimer’s research & therapy 11: 6

23 Chodobski A, Zink BJ, Szmydynger-Chodobska J (2011) Blood–brain barrier pathophysiology in traumatic brain injury. Translational stroke research 2: 492–516

24 Cieri MB, Ramos AJ (2025) Astrocytes, reactive astrogliosis, and glial scar formation in traumatic brain injury. Neural Regeneration Research 20: 973–989

25 Cole JH, Jolly A, de Simoni S, Bourke N, Patel MC, Scott G, Sharp DJ (2018) Spatial patterns of progressive brain volume loss after moderate-severe traumatic brain injury. Brain 141: 822–836 Doi 10.1093/brain/awx354

26 Crupi R, Cordaro M, Cuzzocrea S, Impellizzeri D (2020) Management of traumatic brain injury: from present to future. Antioxidants 9: 297

27 Delgado-García LM, Ojalvo-Sanz AC, Nakamura TK, Martín-López E, Porcionatto M, Lopez-Mascaraque L (2024) Dissecting reactive astrocyte responses: lineage tracing and morphology-based clustering. Biological research 57: 54

28 Early AN, Gorman AA, Van Eldik LJ, Bachstetter AD, Morganti JM (2020) Effects of advanced age upon astrocyte-specific responses to acute traumatic brain injury in mice. Journal of Neuroinflammation 17: 1–16

29 Ekmark-Lewén S, Flygt J, Kiwanuka O, Meyerson BJ, Lewén A, Hillered L, Marklund N (2013) Traumatic axonal injury in the mouse is accompanied by a dynamic inflammatory response, astroglial reactivity and complex behavioral changes. Journal of Neuroinflammation 10: 44

30 Ekmark-Lewén S, Lewén A, Israelsson C, Li GL, Farooque M, Olsson Y, Ebendal T, Hillered L (2010) Vimentin and GFAP responses in astrocytes after contusion trauma to the murine brain. Restorative neurology and neuroscience 28: 311–321

31 El Baassiri MG, Rahal SS, Fulton WB, Sodhi CP, Hackam DJ, Nasr IW (2023) Pharmacologic Toll-like receptor 4 inhibition skews toward a favorable A1/A2 astrocytic ratio improving neurocognitive outcomes following traumatic brain injury. Journal of Trauma and Acute Care Surgery 95: 361–367

32 Endo F, Kasai A, Soto JS, Yu X, Qu Z, Hashimoto H, Gradinaru V, Kawaguchi R, Khakh BS (2022) Molecular basis of astrocyte diversity and morphology across the CNS in health and disease. Science 378: eadc9020

33 Escartin C, Galea E, Lakatos A, O’Callaghan JP, Petzold GC, Serrano-Pozo A, Steinhäuser C, Volterra A, Carmignoto G, Agarwal A (2021) Reactive astrocyte nomenclature, definitions, and future directions. Nature neuroscience 24: 312–325

34 Fesharaki-Zadeh A, Datta D (2024) An overview of preclinical models of traumatic brain injury (TBI): relevance to pathophysiological mechanisms. Frontiers in cellular neuroscience 18: 1371213

35 Fil JE, Joung S, Zimmerman BJ, Sutton BP, Dilger RN (2021) High-resolution magnetic resonance imaging-based atlases for the young and adolescent domesticated pig (Sus scrofa). Journal of Neuroscience Methods 354: 109107

36 Freire MAM, Rocha GS, Bittencourt LO, Falcao D, Lima RR, Cavalcanti JRLP (2023) Cellular and molecular pathophysiology of traumatic brain injury: what have we learned so far? Biology 12: 1139

37 George KK, Heithoff BP, Shandra O, Robel S (2022) Mild traumatic brain injury/concussion initiates an atypical astrocyte response caused by blood–brain barrier dysfunction. Journal of Neurotrauma 39: 211–226

38 Gilchrist J, Thomas K, Wald M, Langlois J (2007) Nonfatal traumatic brain injuries from sports and recreation activities--United States, 2001-2005. MMWR: Morbidity & Mortality Weekly Report 56:

39 Girgis F, Pace J, Sweet J, Miller JP (2016) Hippocampal neurophysiologic changes after mild traumatic brain injury and potential neuromodulation treatment approaches. Frontiers in systems neuroscience 10: 8

40 Graham NS, Jolly A, Zimmerman K, Bourke NJ, Scott G, Cole JH, Schott JM, Sharp DJ (2020) Diffuse axonal injury predicts neurodegeneration after moderate–severe traumatic brain injury. Brain 143: 3685–3698

41 Guskiewicz KM, Marshall SW, Bailes J, McCrea M, Cantu RC, Randolph C, Jordan BD (2005) Association between recurrent concussion and late-life cognitive impairment in retired professional football players. Neurosurgery 57: 719–726

42 Hajiaghamemar M, Margulies SS (2021) Multi-scale white matter tract embedded brain finite element model predicts the location of traumatic diffuse axonal injury. Journal of Neurotrauma 38: 144–157

43 Hajiaghamemar M, Seidi M, Margulies SS (2020) Head rotational kinematics, tissue deformations, and their relationships to the acute traumatic axonal injury. Journal of biomechanical engineering 142: 031006

44 Hajiaghamemar M, Seidi M, Oeur RA, Margulies SS (2019) Toward development of clinically translatable diagnostic and prognostic metrics of traumatic brain injury using animal models: A review and a look forward. Experimental neurology 318: 101–123

45 Hajiaghamemar M, Wu T, Panzer MB, Margulies SS (2020) Embedded axonal fiber tracts improve finite element model predictions of traumatic brain injury. Biomechanics and modeling in mechanobiology 19: 1109–1130

46 Hirrlinger J, Nimmerjahn A (2022) A perspective on astrocyte regulation of neural circuit function and animal behavior. Glia 70: 1554–1580

47 Huang X-j, Glushakova O, Mondello S, Van K, Hayes RL, Lyeth BG (2015) Acute temporal profiles of serum levels of UCH-L1 and GFAP and relationships to neuronal and astroglial pathology following traumatic brain injury in rats. Journal of neurotrauma 32: 1179–1189

48 Hutchinson EB, Romero-Lozano A, Johnson HR, Knutsen AK, Bosomtwi A, Korotcov A, Shunmugavel A, King SG, Schwerin SC, Juliano SL (2022) Translationally Relevant Magnetic Resonance Imaging Markers in a Ferret Model of Closed Head Injury. Frontiers in Neuroscience 15: 779533

49 Hutchinson EB, Schwerin SC, Radomski K, Sadeghi N, Jenkins J, Komlosh ME, Irfanoglu MO, Juliano SL, Pierpaoli C (2017) Population based MRI and DTI templates of the adult ferret brain and tools for voxelwise analysis. Neuroimage 152: 575–589

50 Ichkova A, Badaut J (2017) New biomarker stars for traumatic brain injury. Journal of Cerebral Blood Flow & Metabolism 37: 3276–3277

51 Jafari B, Memar M (2025) Exploring Mesoscale Brain Connectivity Variations and Developing Sex-Specific Tractography Templates. NeuroImage: 121545

52 Johnson GA, Badea A, Brandenburg J, Cofer G, Fubara B, Liu S, Nissanov J (2010) Waxholm space: an image-based reference for coordinating mouse brain research. Neuroimage 53: 365–372

53 Johnson GA, Laoprasert R, Anderson RJ, Cofer G, Cook J, Pratson F, White LE (2021) A multicontrast MR atlas of the Wistar rat brain. NeuroImage 242: 118470

54 Johnson VE, Stewart W, Smith DH (2013) Axonal pathology in traumatic brain injury. Experimental neurology 246: 35–43

55 Jurga AM, Paleczna M, Kadluczka J, Kuter KZ (2021) Beyond the GFAP-astrocyte protein markers in the brain. Biomolecules 11: 1361

56 Kahriman A, Bouley J, Smith TW, Bosco DA, Woerman AL, Henninger N (2021) Mouse closed head traumatic brain injury replicates the histological tau pathology pattern of human disease: characterization of a novel model and systematic review of the literature. Acta Neuropathologica Communications 9: 118

57 Karve IP, Taylor JM, Crack PJ (2016) The contribution of astrocytes and microglia to traumatic brain injury. British journal of pharmacology 173: 692–702

58 Kawata K, Rubin LH, Lee JH, Sim T, Takahagi M, Szwanki V, Bellamy A, Darvish K, Assari S, Henderer JD (2016) Association of football subconcussive head impacts with ocular near point of convergence. JAMA ophthalmology 134: 763–769

59 Kayasandik CB, Ru W, Labate D (2020) A multistep deep learning framework for the automated detection and segmentation of astrocytes in fluorescent images of brain tissue. Scientific reports 10: 5137

60 Khatri N, Sumadhura B, Kumar S, Kaundal RK, Sharma S, Datusalia AK (2021) The complexity of secondary cascade consequent to traumatic brain injury: pathobiology and potential treatments. Current Neuropharmacology 19: 1984

61 Khodadadei F, Arshad R, Morales DM, Gluski J, Marupudi NI, McAllister JP, Limbrick Jr DD, Harris CA (2022) The effect of A1 and A2 reactive astrocyte expression on hydrocephalus shunt failure. Fluids and Barriers of the CNS 19: 78

62 Krieg JL, Leonard AV, Tuner RJ, Corrigan F (2023) Characterization of traumatic brain injury in a gyrencephalic ferret model using the novel closed head injury model of engineered rotational acceleration (CHIMERA). Neurotrauma Reports 4: 761–780

63 Krieg JL, Leonard AV, Turner RJ, Corrigan F (2023) Identifying the Phenotypes of Diffuse Axonal Injury Following Traumatic Brain Injury. Brain sciences 13: 1607

64 Kugler E, Bravo I, Durmishi X, Marcotti S, Beqiri S, Carrington A, Stramer B, Mattar P, MacDonald RB (2023) GliaMorph: A modular image analysis toolkit to quantify Müller glial cell morphology. Development 150: dev201008

65 Kuo GY, Tarzi FP, Louie S, Poblete RA (2022) Neuroinflammation in Traumatic Brain Injury. Frontiers In Traumatic Brain Injury. IntechOpen, City

66 Lattke M, Guillemot F (2022) Understanding astrocyte differentiation: Clinical relevance, technical challenges, and new opportunities in the omics era. WIREs Mechanisms of Disease 14: e1557

67 Lawrence JM, Schardien K, Wigdahl B, Nonnemacher MR (2023) Roles of neuropathology-associated reactive astrocytes: a systematic review. Acta neuropathologica communications 11: 42

68 Lecky FE, Otesile O, Marincowitz C, Majdan M, Nieboer D, Lingsma HF, Maegele M, Citerio G, Stocchetti N, Steyerberg EW (2021) The burden of traumatic brain injury from low-energy falls among patients from 18 countries in the CENTER-TBI Registry: A comparative cohort study. PLoS medicine 18: e1003761

69 Lee AL (2020) Advanced imaging of traumatic brain injury. Korean Journal of Neurotrauma 16: 3

70 Lee H-H, Park S-C, Choe I-S, Kim Y, Ha Y-S (2015) Time Course and Characteristics of Astrocyte Activation in the Rat Brain after Injury. Korean journal of neurotrauma 11: 44–51

71 Li D-R, Zhang F, Wang Y, Tan X-H, Qiao D-F, Wang H-J, Michiue T, Maeda H (2012) Quantitative analysis of GFAP-and S100 protein-immunopositive astrocytes to investigate the severity of traumatic brain injury. Legal medicine 14: 84–92

72 Liu J, Kou Z, Tian Y (2014) Diffuse axonal injury after traumatic cerebral microbleeds: an evaluation of imaging techniques. Neural regeneration research 9: 1222–1230

73 Liu L-r, Liu J-c, Bao J-s, Bai Q-q, Wang G-q (2020) Interaction of microglia and astrocytes in the neurovascular unit. Frontiers in immunology 11: 1024

74 McAteer KM, Corrigan F, Thornton E, Turner RJ, Vink R (2016) Short and long term behavioral and pathological changes in a novel rodent model of repetitive mild traumatic brain injury. PLoS One 11: e0160220

75 Mckee AC, Daneshvar DH (2015) The neuropathology of traumatic brain injury. Handbook of clinical neurology 127: 45–66

76 Michinaga S, Koyama Y (2021) Pathophysiological responses and roles of astrocytes in traumatic brain injury. International journal of molecular sciences 22: 6418

77 Mira RG, Lira M, Cerpa W (2021) Traumatic brain injury: mechanisms of glial response. Frontiers in physiology 12: 740939

78 Moulson AJ, Squair JW, Franklin RJ, Tetzlaff W, Assinck P (2021) Diversity of reactive astrogliosis in CNS pathology: heterogeneity or plasticity? Frontiers in cellular neuroscience 15: 703810

79 Mouzon B, Chaytow H, Crynen G, Bachmeier C, Stewart J, Mullan M, Stewart W, Crawford F (2012) Repetitive mild traumatic brain injury in a mouse model produces learning and memory deficits accompanied by histological changes. Journal of neurotrauma 29: 2761–2773

80 Muñoz-Ballester C, Robel S (2023) Astrocyte-mediated mechanisms contribute to traumatic brain injury pathology. WIREs Mechanisms of Disease 15: e1622

81 O’Connor WT, Smyth A, Gilchrist MD (2011) Animal models of traumatic brain injury: a critical evaluation. Pharmacology & therapeutics 130: 106–113

82 Ohtake Y, Smith GM, Li S (2016) Reactive astrocyte scar and axon regeneration: suppressor or facilitator? Neural Regeneration Research 11: 1050–1051

83 Özen I, Mai H, De Maio A, Ruscher K, Michalettos G, Clausen F, Gottschalk M, Ansar S, Arkan S, Erturk A (2022) Purkinje cell vulnerability induced by diffuse traumatic brain injury is linked to disruption of long-range neuronal circuits. Acta neuropathologica communications 10: 129

84 Papa L, Brophy GM, Welch RD, Lewis LM, Braga CF, Tan CN, Ameli NJ, Lopez MA, Haeussler CA, Mendez Giordano DI (2016) Time course and diagnostic accuracy of glial and neuronal blood biomarkers GFAP and UCH-L1 in a large cohort of trauma patients with and without mild traumatic brain injury. JAMA neurology 73: 551–560

85 Pekny M, Pekna M (2014) Astrocyte reactivity and reactive astrogliosis: costs and benefits. Physiological reviews 94: 1077–1098

86 Peterson A, Thomas K, Zhou H (2018) Traumatic brain injury-related deaths by age group, sex, and mechanism of injury. CDC TBI surveillance report:

87 Quesenberry KE, de Matos R (2020) Basic approach to veterinary care of ferrets. Ferrets, rabbits, and rodents: 13

88 Ramzanpour M, Jafari B, Smith J, Allen J, Hajiaghamemar M (2023) Comprehensive study of sex-based anatomical variations of human brain and development of sex-specific brain templates. Brain Multiphysics 4: 100077

89 Roboon J, Hattori T, Nguyen DT, Ishii H, Takarada-Iemata M, Kannon T, Hosomichi K, Maejima T, Saito K, Shinmyo Y (2022) Isolation of ferret astrocytes reveals their morphological, transcriptional, and functional differences from mouse astrocytes. Frontiers in Cellular Neuroscience 16: 877131

90 Rui Q, Ni H, Lin X, Zhu X, Li D, Liu H, Chen G (2019) Astrocyte-derived fatty acid-binding protein 7 protects blood-brain barrier integrity through a caveolin-1/MMP signaling pathway following traumatic brain injury. Experimental neurology 322: 113044

91 Scale NS National Institute of Neurological Disorders and Stroke. 2020. City

92 Schachtrup C, Ryu JK, Helmrick MJ, Vagena E, Galanakis DK, Degen JL, Margolis RU, Akassoglou K (2010) Fibrinogen triggers astrocyte scar formation by promoting the availability of active TGF-â after vascular damage. Journal of Neuroscience 30: 5843–5854

93 Schwerin SC, Chatterjee M, Hutchinson EB, Djankpa FT, Armstrong RC, McCabe JT, Perl DP, Juliano SL (2021) Expression of GFAP and tau following blast exposure in the cerebral cortex of ferrets. Journal of Neuropathology & Experimental Neurology 80: 112–128

94 Schwerin SC, Chatterjee M, Imam-Fulani AO, Radomski KL, Hutchinson EB, Pierpaoli CM, Juliano SL (2018) Progression of histopathological and behavioral abnormalities following mild traumatic brain injury in the male ferret. Journal of neuroscience research 96: 556–572

95 Schwerin SC, Hutchinson EB, Radomski KL, Ngalula KP, Pierpaoli CM, Juliano SL (2017) Establishing the ferret as a gyrencephalic animal model of traumatic brain injury: optimization of controlled cortical impact procedures. Journal of neuroscience methods 285: 82–96

96 Sethi P, Virmani G, Gupta K, Thumu SCR, Ramanan N, Marathe S (2021) Automated morphometric analysis with SMorph software reveals plasticity induced by antidepressant therapy in hippocampal astrocytes. Journal of cell science 134: jcs258430

97 Shandra O, Winemiller AR, Heithoff BP, Munoz-Ballester C, George KK, Benko MJ, Zuidhoek IA, Besser MN, Curley DE, Edwards III GF (2019) Repetitive diffuse mild traumatic brain injury causes an atypical astrocyte response and spontaneous recurrent seizures. The Journal of Neuroscience 39: 1944–1963

98 SheikhBahaei S, Morris B, Collina J, Anjum S, Znati S, Gamarra J, Zhang R, Gourine AV, Smith JC (2018) Morphometric analysis of astrocytes in brainstem respiratory regions. Journal of Comparative Neurology 526: 2032–2047

99 Shlosberg D, Benifla M, Kaufer D, Friedman A (2010) Blood–brain barrier breakdown as a therapeutic target in traumatic brain injury. Nature Reviews Neurology 6: 393–403

100 Silvestro S, Raffaele I, Quartarone A, Mazzon E (2024) Innovative insights into traumatic brain injuries: biomarkers and new pharmacological targets. International Journal of Molecular Sciences 25: 2372

101 Smith DH, Kochanek PM, Rosi S, Meyer R, Ferland-Beckham C, Prager EM, Ahlers ST, Crawford F (2021) Roadmap for advancing pre-clinical science in traumatic brain injury. Journal of Neurotrauma 38: 3204–3221

102 Smith DH, Meaney DF, Shull WH (2003) Diffuse axonal injury in head trauma. The Journal of head trauma rehabilitation 18: 307–316

103 Sofroniew MV (2015) Astrogliosis. Cold Spring Harbor perspectives in biology 7: a020420

104 Sofroniew MV (2009) Molecular dissection of reactive astrogliosis and glial scar formation. Trends in neurosciences 32: 638–647

105 Sofroniew MV, Vinters HV (2010) Astrocytes: biology and pathology. Acta neuropathologica 119: 7–35

106 Sun D, Lye-Barthel M, Masland RH, Jakobs TC (2010) Structural remodeling of fibrous astrocytes after axonal injury. Journal of Neuroscience 30: 14008–14019

107 Tavares G, Martins M, Correia JS, Sardinha VM, Guerra-Gomes S, das Neves SP, Marques F, Sousa N, Oliveira JF (2017) Employing an open-source tool to assess astrocyte tridimensional structure. Brain Structure and Function 222: 1989-1999

108 Thomas BP, Liu P, Park DC, Van Osch MJ, Lu H (2014) Cerebrovascular reactivity in the brain white matter: magnitude, temporal characteristics, and age effects. Journal of Cerebral Blood Flow & Metabolism 34: 242–247

109 Villapol S, Byrnes KR, Symes AJ (2014) Temporal dynamics of cerebral blood flow, cortical damage, apoptosis, astrocyte–vasculature interaction and astrogliosis in the pericontusional region after traumatic brain injury. Frontiers in neurology 5: 82

110 Wang N, Anderson RJ, Ashbrook DG, Gopalakrishnan V, Park Y, Priebe CE, Qi Y, Laoprasert R, Vogelstein JT, Williams RW (2020) Variability and heritability of mouse brain structure: Microscopic MRI atlases and connectomes for diverse strains. Neuroimage 222: 117274

111 Wang N, Anderson RJ, Badea A, Cofer G, Dibb R, Qi Y, Johnson GA (2018) Whole mouse brain structural connectomics using magnetic resonance histology. Brain Structure and Function 223: 4323–4335

112 Weeks D, Sullivan S, Kilbaugh T, Smith C, Margulies SS (2014) Influences of developmental age on the resolution of diffuse traumatic intracranial hemorrhage and axonal injury. Journal of neurotrauma 31: 206–214

113 Wei DC, Morrison EH (2019) Histology, Astrocytes.

114 Wilhelmsson U, Bushong EA, Price DL, Smarr BL, Phung V, Terada M, Ellisman MH, Pekny M (2006) Redefining the concept of reactive astrocytes as cells that remain within their unique domains upon reaction to injury. Proceedings of the National Academy of Sciences 103: 17513–17518

115 Yang Z, Wang KK (2015) Glial fibrillary acidic protein: from intermediate filament assembly and gliosis to neurobiomarker. Trends in neurosciences 38: 364–374

116 Zhang H, Zhang X, Chai Y, Wang Y, Zhang J, Chen X (2025) Astrocyte-mediated inflammatory responses in traumatic brain injury: Mechanisms and potential interventions. Frontiers in Immunology 16: 1584577

117 Zhao Q, Zhang J, Li H, Li H, Xie F (2023) Models of traumatic brain injury-highlights and drawbacks. Frontiers in neurology 14: 1151660

118 Zhou Y, Shao A, Yao Y, Tu S, Deng Y, Zhang J (2020) Dual roles of astrocytes in plasticity and reconstruction after traumatic brain injury. Cell Communication and Signaling 18: 1–16

119 Zwirner J, Lier J, Franke H, Hammer N, Matschke J, Trautz F, Tse R, Ondruschka B (2021) GFAP positivity in neurons following traumatic brain injuries. International Journal of Legal Medicine 135: 2323–2333

