## Supplementary Table 1 for "Whole-Brain, Region-Specific Astrocyte Reactivity and Morphological Remodeling After Diffuse Traumatic Brain Injury in A Gyrencephalic Ferret Model"

**Table S1-** Regional fractional volumes of key neuroanatomical regions across species. For cross-species comparison, established MRI atlases from humans (mixed-sex population template; 609 males, 721 females)[1,2] rhesus macaques (10 adults, 2–11 years; 7 males, 3 females)[3], pigs (20 males, 4 weeks)[4], ferrets (26 in vivo and 8 ex vivo scans, 5–10 months)[5], rats (6 adult male Wistar)[6], and mice (14 males, 8 weeks old)[7-9] were used.

| | Human | Primate | Pig | Ferret | Rat | Mouse |
|---|---|---|---|---|---|---|
| CorticalGM | 39.30% | 47.46% | | 42.35% | 36.12% | 27.88% |
| GrayMatter | 11.93% | 12.81% | 22.49% | 17.98% | 20.90% | 22.73% |
| NotMapped | | 2.32% | | | 5.75% | 9.02% |
| SubCorticalGM | 4.46% | 11.34% | 9.95% | 13.90% | 26.72% | 28.79% |
| Sulcus | | | | 3.89% | | |
| WhiteMatter | 42.78% | 27.37% | 37.40% | 21.48% | 10.51% | 9.14% |

| | Human | Primate | Pig | Ferret | Rat | Mouse |
|---|---|---|---|---|---|---|
| Amygdala | 0.24% | 0.65% | 0.02% | 0.25% | 2.20% | 2.96% |
| Brainstem | 2.12% | 3.19% | 5.73% | 5.58% | 9.79% | 6.28% |
| Caudate | 0.70% | 1.66% | 0.85% | 1.88% | | |
| Cerebellum | 11.54% | 9.54% | 0.66% | 14.74% | 15.16% | 10.24% |
| Cingulum | | | | 0.45% | 0.29% | 0.15% |
| CorpusCallosum | 0.22% | | 0.66% | 1.09% | 3.56% | 1.54% |
| CortexWM | 39.90% | 25.46% | 37.40% | 10.34% | | |
| Fornix | | | | 0.97% | 0.77% | 0.60% |
| Hippocampus | 0.73% | 1.08% | 1.28% | 3.63% | 7.39% | 5.93% |
| Hypothalamus | 0.78% | 0.43% | 0.24% | 1.23% | 2.47% | 2.85% |
| Putamen | 0.97% | 2.16% | 5.18% | 0.21% | 4.74% | 5.68% |
| Thalamus | 1.38% | 2.91% | 1.79% | 3.51% | 4.24% | 4.72% |

### References:


1. Ramzanpour M, Jafari B, Smith J, et al. Comprehensive study of sex-based anatomical variations of human brain and development of sex-specific brain templates. Brain Multiphysics 2023;4(100077
2. Jafari B, Memar M. Exploring Mesoscale Brain Connectivity Variations and Developing Sex-Specific Tractography Templates. NeuroImage 2025;121545
3. Calabrese E, Badea A, Coe CL, et al. A diffusion tensor MRI atlas of the postmortem rhesus macaque brain. Neuroimage 2015;117(408-416

4. Fil JE, Joung S, Zimmerman BJ, et al. High-resolution magnetic resonance imaging-based atlases for the young and adolescent domesticated pig (Sus scrofa). Journal of Neuroscience Methods 2021;354(109107
5. Hutchinson EB, Schwerin SC, Radomski K, et al. Population based MRI and DTI templates of the adult ferret brain and tools for voxelwise analysis. Neuroimage 2017;152(575-589
6. Johnson GA, Laoprasert R, Anderson RJ, et al. A multicontrast MR atlas of the Wistar rat brain. NeuroImage 2021;242(118470
7. Johnson GA, Badea A, Brandenburg J, et al. Waxholm space: an image-based reference for coordinating mouse brain research. Neuroimage 2010;53(2):365-372
8. Wang N, Anderson RJ, Badea A, et al. Whole mouse brain structural connectomics using magnetic resonance histology. Brain Structure and Function 2018;223(9):4323-4335
9. Wang N, Anderson RJ, Ashbrook DG, et al. Variability and heritability of mouse brain structure: Microscopic MRI atlases and connectomes for diverse strains. Neuroimage 2020;222(117274
